# Bacterial cell cycle and growth phase switch by the essential transcriptional regulator CtrA

**DOI:** 10.1101/739516

**Authors:** Marie Delaby, Gaël Panis, Patrick H. Viollier

## Abstract

Many bacteria acquire dissemination and virulence traits in G1-phase. CtrA, an essential and conserved cell cycle transcriptional regulator identified in the dimorphic alpha-proteobacterium *Caulobacter crescentus*, first activates promoters in late S-phase and then mysteriously switches to different target promoters in G1-phase. We uncovered a highly conserved determinant in the DNA-binding domain (DBD) of CtrA uncoupling this promoter switch. We also show that it reprograms CtrA occupancy in stationary cells inducing a (p)ppGpp alarmone signal perceived by the RNA polymerase beta subunit. A simple side chain modification in a critical residue within the core DBD imposes opposing developmental phenotypes and transcriptional activities of CtrA. A naturally occurring polymorphism in the rickettsial DBD resembles a mutation that drives CtrA towards activation of the dispersal (G1-phase) program in *Caulobacter*. Hence, we propose that this determinant dictates promoter reprogramming during the growth transition of obligate intracellular rickettsia differentiating from replicative cells into dispersal cells.

## Introduction

Tight regulation of gene expression during the cell cycle is paramount to ensure proper timing and coordination of DNA replication, chromosome segregation and cell division, often concurrently with morphological changes. *Caulobacter crescentus* is a rod-shaped and dimorphic alpha-proteobacterium that undergoes an asymmetric cell division into two unequally sized and polarized daughter cells: a motile and non-replicative dispersal (swarmer, SW) cell residing in G1-phase and a capsulated and replicative (stalked, ST) cell. In *C. crescentus,* cell cycle progression is intimately tied to polar remodeling via a circuit of transcriptional activators that direct sequential gene expression programs^1^. The G1-phase program is implemented in the SW daughter cell that inherits the new cell pole where the flagellum and adhesive pili are located. By contrast, the old pole is inherited by the replicative ST cell that is engaged in DNA-replication. As the cell cycle proceeds, the ST cell prepares for division, expresses the late S-phase program and polarizes before dividing asymmetrically into a SW and ST daughter cell (Figure 1A). The transcriptional programs are not only temporally ordered, but also spatially confined during cytokinesis, with the G1-phase program being activated in the nascent SW chamber during cytokinesis, but not in the ST cell chamber^1, 2^.

**Figure 1.**
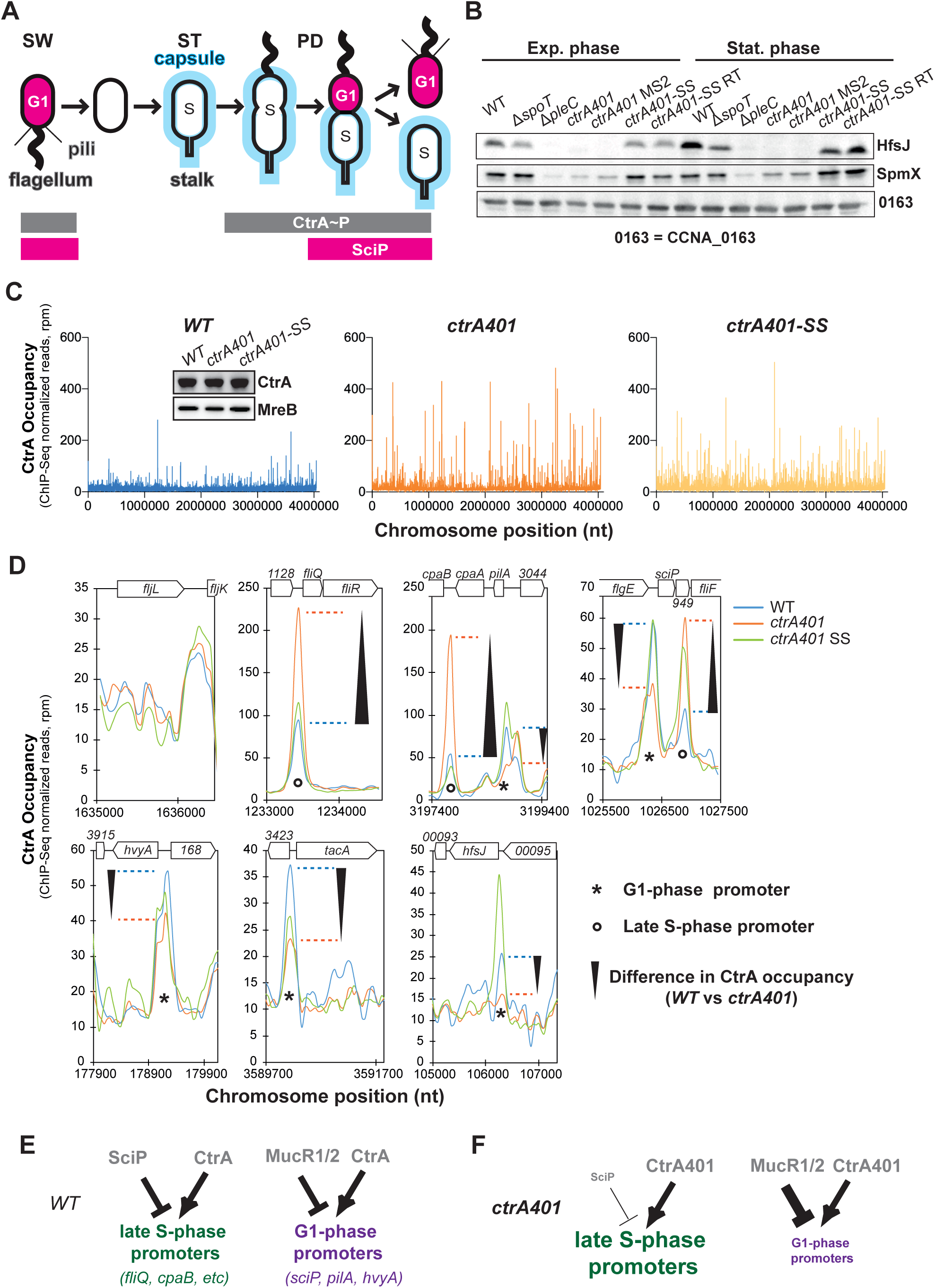
*ctrA401* as gain of function mutation. A) Schematic of *Caulobacter* capsulation (blue) along cell cycle and regulatory interactions that controls *Caulobacter* transcriptional switch between late S-phase and G1-phase. CtrA controls activity of late S- and G1-phase promoters. SciP which is expressed in G1-phase under the control of CtrA negatively regulates S-phase genes. During cell cycle, capsulation is regulated through expression of *hvyA* that prevents capsulation in Sw cell and under the control of the transcriptional regulators CtrA. B) Immunoblots showing steady-state levels of HfsJ and SpmX in WT, Δ*spoT*, Δ*pleC*, *ctrA401* and derivatives in exponential and stationary phase. CCNA_00163 serves as a loading control. C) Genome wide occupancies of CtrA on the Caulobacter *WT*, *ctrA401* and *ctrA401-SS* genome as determined by ChIP-seq. The x-axis represents the nucleotide position on the genome (bp), whereas the y-axis shows the normalized ChIP profiles in read per million (rpm). D) ChIP-seq traces of CtrA, CtrA401 (T170I) and CtrA401-SS (T168I/T170I) on different CtrA target promoters. Genes encoded are represented as boxes on the upper part of the graph, gene names and CCNA numbers gene annotation are indicated in the boxes or above. E, F) Schemes showing the regulatory interactions happening at the late S- and G-phase promoters based on C, D and Table 1.

Cell cycle analyses are facile with *C. crescentus* because the non-capsulated G1-phase (SW) cells can be separated from capsulated S-phase (ST) cells by density gradient centrifugation^3^. Replicative functions are acquired with the obligate G1➔S-phase transition that morphologically manifests with the differentiation from SW to ST cells. Pili and the flagellum are lost from the same cell pole, followed by the onset of stalk outgrowth from the vacated site^4^. Concurrently, the capsule is synthesized which increases the cellular buoyancy^5^, and DNA synthesis initiates bidirectionally from a single origin of replication (*Cori,* Figure 1A) on the circular chromosome. A multicomponent genetic circuit of global transcriptional activators and repressors temporally and spatially coordinates these events at the molecular level^1^.

The late S-phase and G1-phase programs are both activated by CtrA, a highly conserved member of the OmpR family of DNA-binding response regulators^1, 2^. CtrA is essential for viability and proper cell cycle control in *C. crescentus*^6^ and in many other alpha-proteobacteria^1^. CtrA switches from activating the late S-phase promoters before cell division to inducing G1-phase promoters in the nascent SW cell chamber at cytokinesis. While CtrA also binds *Cori* and prevents the initiation of DNA replication in G1-phase^6–8^, it is degraded by the ClpXP protease during the G1➔S transition^9^. It is re-synthesized in late S-phase and again degraded in the ST compartment during cytokinesis, while being maintained in the SW compartment (Figure 1A). The conserved target motif (CtrA box: 5’-TTAA-N7-TTAA-3’) is present in both promoter classes and recognized by the C-terminal DNA binding domain (DBD) of CtrA. At the N-terminus, CtrA harbors a receiver domain (RD) with a phosphorylation site at a conserved aspartate (at position 51, D51). Phosphorylation at D51 stimulates DNA binding and is required for viability. The hybrid histidine kinase CckA directs a multi-component phosphoryl-transfer reaction to D51 of CtrA^10, 11^. Though loss of CckA is lethal, missense mutations in the RD were isolated in unbiased selection for mutant CtrA derivatives that can support viability of *C. crescentus* cells lacking CckA^12^.

Mutations in the DBD domain of CtrA that are critical for viability have also been isolated^6^. In the landmark study by Quon *et al*, *ctrA* was uncovered as an essential gene in *C. crescentus* [as the *ctrA401* mutant allele, encoding CtrA(T170I)] in a two-step genetic selection. First, based on earlier evidence that the *fliQ* (class II) flagellar assembly gene is transcriptionally de-repressed in late S-phase, the authors selected for mutants with elevated *fliQ* promoter (P*_fliQ_*) activity at 28°C. Next, those mutants were retained and those that also exhibit thermosensitive growth at 37°C compared to wild-type (*WT*), yielding the *ctrA401* mutant. Since P*_fliQ_* activity is elevated at 28°C, but strongly impaired at 37°C in *ctrA401* cells, it was concluded that CtrA acts positively and negatively at P*_fliQ_* (and likely other late S-phase promoters)^6^.

How CtrA switches its specificity from late S-phase promoters to G1-phase promoters is unclear. Determinants in CtrA that are specific for each promoter class have not been identified. At least two different negative regulators, one targeting the late S-phase promoters and another acting on G1-phase promoters^12–14^, reinforce the promoter switch^12^. The conserved helix-turn-helix protein SciP specifically inhibits late S-phase promoters that are activated by CtrA^12, 15, 16^. SciP is restricted to G1-phase due in part to its synthesis from a CtrA-activated promoter (P*_sciP_*) that fires in G1-phase and in part due to a short half-life imparted by the Lon protease^9, 12, 15, 16^. Thus, SciP couples the activation of G1-phase promoters with the shutdown of late S-phase promoter activity, but it cannot act as the trigger for the promoter switch. The repressor paralogs MucR1 and MucR2 negatively regulate the CtrA-activated promoters that fire in G1-phase. While MucR1/2 abundance and (steady-state) promoter occupancy does not change during the cell cycle, the ability of MucR1/2 to exclude competing proteins from target promoters is reduced in G1-phase^17^. Interestingly, MucR orthologs regulate virulence gene expression in other alpha-proteobacteria and bind orthologous promoters^12^. These free-living alpha-proteobacteria express virulence traits in G1-phase, suggesting that the mechanism of restricting promoter firing to G1-phase by MucR is conserved^12^.

In addition to its cell cycle function, CtrA has been implicated in regulating development during a growth phase transition in obligate intracellular alpha-proteobacteria from the order Rickettsiales^18^. Akin to the developmental cycle in other obligate intracellular bacteria, the dimorphic human pathogen *Ehrlichia chaffeensis* differentiates into infectious dispersal (dense-cored) cells following the rapid intracellular expansion of the replicative (reticulate) cells^19^, suggesting a nutritional trigger may underlie development. Rickettsia do not encode MucR or SciP orthologs in their genomes^1^, but CtrA peaks during the late stages of growth when the transition from reticulate to dense-cored cells occurs^18^. In stationary phase, a starvation stress response is triggered in *C. crescentus* cells by the alarmone (p)ppGpp (guanosine tetraphosphate and guanosine pentaphosphate) that supresses DNA replication and maintains CtrA abundance to arrest cells predominantly in G1 phase and to a lesser extent in the predivisional phase^20^. Whether CtrA is also involved in transcriptional re-programming during this growth phase transition in *C. crescentus* cells has not been investigated.

Here we unearth a molecular determinant within the DBD of CtrA that is required to execute the switch from late S- to G1- phase promoters and to reprogram CtrA in stationary *C. crescentus* cells accumulating (p)ppGpp. A highly conserved triad centered in the core DBD dictates whether CtrA can switch G1-phase promoters and use it to control the cell cycle during a growth phase transition induced by (p)ppGpp in response to starvation stress. CtrA activity can be maintained in cells expressing a mutant form of RNA polymerase that is blind to (p)ppGpp, indicating that transcriptional regulation coordinates growth phase transition regulated by CtrA.

## Materiel and Methods

### Growth conditions

*Caulobacter crescentus* NA1000 and derivatives were grown at 30°C in PYE (peptone yeast extract). Antibiotic concentrations used for *C. crescentus* include kanamycin (solid: 20 μg/ml; liquid: 5 μg/ml), tetracycline (1 μg/ml), gentamycin (1 μg/ml) and nalidixic acid (20 μg/ml). When needed, D-xylose or sucrose was added at 0.3% final concentration, glucose at 0.2% final concentration and vanillate at 500 μM final concentration. *Escherichia coli* S17-1 λ*pir* and EC100D (Epicentre Technologies, Madison, WI, USA) were grown at 37°C in LB. Swarmer cell isolation, electroporations, biparental mattings and bacteriophage ϕCr30-mediated generalized transductions were performed as previously described^3^. Plasmids for β -galactosidase assays were into *C. crescentus* by electroporation.

### Bacterial Strains, Plasmids, and Oligonucleotides

Bacterial strains, plasmids, and oligonucleotides used in this study are listed and described in supplementary material and methods.

### Motility suppressors of *ctrA401* mutant cells

Spontaneous mutations that suppress the motility defect of the *ctrA401* mutant cells appeared as “flares” that emanated from nonmotile colonies after approximately 4 to 5 days of incubation at 30°C. Ten isolates (MS 1 to 10) were selected and tested for the excision of the MGE by PCR using primers amplifying the *CCNA_00477* (presence of the MGE) and *attB* (MGE excised) fragments. Two *ctrA401* MS (MS 2 and 4) were subjected to whole genome sequencing. The Nextera kit from illumina was used for library preparation with 50 ng of DNA as input. The Library molarity and quality was assessed with the Qubit and Tapestation using a DNA High sensitivity chip (Agilent Technologies). Libraries were loaded on a HiSeq 4000 single-read Illumina flow cell. Reads of 50 bases were generated. The nine other suppressors were then sequenced by Sanger sequencing for the *CCNA_03998* (amplified using primers 3998_NdeI and 3998_EcoRI). The table 1 below summarizes the mutations obtained in the suppressors. To confirm that these mutations were responsible for the suppression of the motility defect of the *ctrA401* mutant cells, ΔMGE *ctrA401* and Δ*3998 ctrA401* double mutants were constructed through ϕCr30 mediated generalized transduction of the *ctrA401* allele. The transducing phage stock is a lysate of *ctrA401* MS2 *ctrA401::*pNPTS138*-ctrA-ds*. Sucrose 3% was then used to eliminate the pNPTS138 by homologous recombination and clones were screen for kanamycin susceptibility. Sanger sequencing was used to verify the integrity of the mutation using primers ctrA_NdeI and ctrA_XbaI.

**Table 1.**
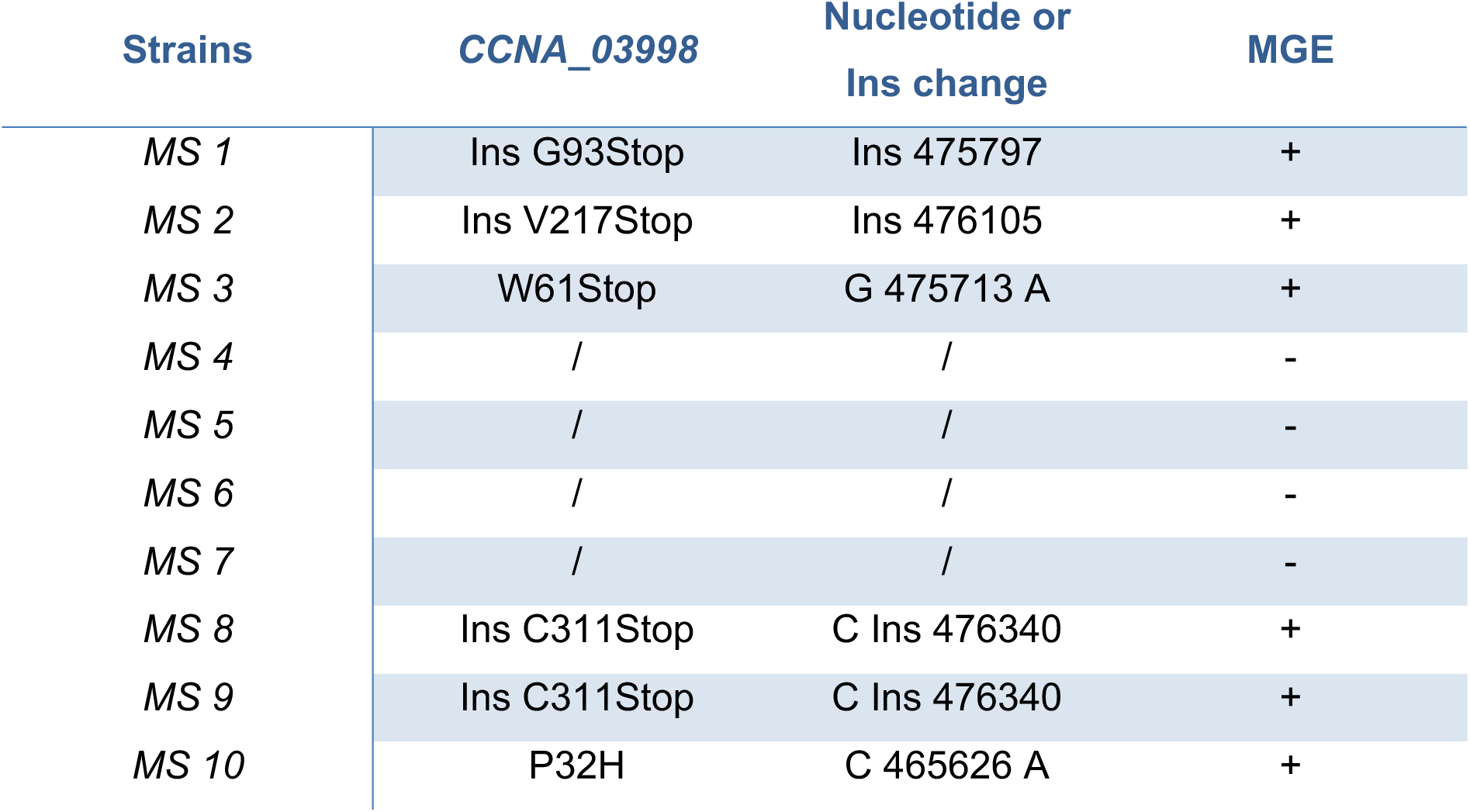
Mutations identified in the ctrA401 <u>M</u>otility <u>S</u>uppressors.

| Strains | CCNA_03998 | Nucleotide or<br>Ins change | MGE |
| --- | --- | --- | --- |
| MS 1 | Ins G93Stop | Ins 475797 | + |
| MS 2 | Ins V217Stop | Ins 476105 | + |
| MS 3 | W61Stop | G 475713 A | + |
| MS 4 | / | / | - |
| MS 5 | / | / | - |
| MS 6 | / | / | - |
| MS 7 | / | / | - |
| MS 8 | Ins C311Stop | C Ins 476340 | + |
| MS 9 | Ins C311Stop | C Ins 476340 | + |
| MS 10 | P32H | C 465626 A | + |

### Motility suppressors of NA1000 Δ*ptsP* Δ*spoT* mutant cells

Spontaneous mutations that suppress the motility defect of the Δ*ptsP* Δ*spoT* mutant cell appeared as “flares” that emanated from nonmotile colonies after approximately 4 to 5 days of incubation at 30°C. One isolate (UG5591) was subjected to whole genome sequencing, and mutation in the *rpoB* (*rpoB^H559Y^*) gene was found. In the suppressor, the histidine codon at position 559 in *rpoB* was changed to one encoding tyrosine (change of nt 549505 from C to T relative to the NA1000 genome sequence). To confirm that this mutation was responsible for improving the motility defect of the Δ*ptsP* Δ*spoT* mutant cells, we backcross the mutation *rpoB^H559Y^* into *WT* cells. To this end, a pNPTS138-*rpoC*-ds, conferring kanamycin resistance, was integrated by homologous recombination downstream of the *rpoC* gene in the Δ*ptsP* Δ*spoT rpoB^H559Y^* mutant. Next, ϕCr30 mediated generalized transduction was used to transfer the *rpoB^H559Y^* allele into the *WT* cells. Sucrose 3% was then used to eliminate the pNPTS138-*rpoC*-ds by homologous recombination and clones were screen for kanamycin susceptibility. Sanger sequencing was used to verify the integrity of the mutation using primers rpoB-seq1 and rpoB-seq2.

### ϕCr30-mediated transduction of the *ctrA401* allele

*ctrA401* transducing phage stock is a lysate of *ctrA401* MS2 *ctrA401::*pNPTS138*-ctrA-ds*.

### Genome-wide transposon mutagenesis coupled to deep sequencing (Tn-Seq)

#### NA1000 ctrA401 MS2 pilA::P_pilA_-nptII Tn-Seq

Transposon mutagenesis of *C. crescentus* NA1000 *ctrA401* MS2 *pilA*::P*_pilA_*-*nptII* was done by intergeneric conjugation from *E. coli* S17-1 λ*pir* harbouring the *himar1*-derivative p*MAR2xT7*^21^. For the *C. crescentus* NA1000 *ctrA401* MS2 *pilA*::P*_pilA_*-*nptII*::Tn, a Tn-library of >200,000 gentamicin- and kanamycin-resistant clones was collected. NA1000 *ctrA401* MS2 *pilA*::P*_pilA_*-*nptII*::Tn(Gent^R^) bank was grown overnight in PYE and chromosomal DNA was extracted. Genomic DNA was used to generate barcoded Tn-Seq libraries and submitted to Illumina HiSeq 4000 sequencing (Fasteris SA). Tn insertion-specific reads (150 bp long) were sequenced using the himar-Tnseq2 primer (5’-AGACCGGGGACTTATCAGCCAACCTGT-3’). Specific reads attesting an integration of the transposon on a 5’-TA-3’ specific DNA locus were sorted (Rstudio_V1.1.442) from the tens of million reads generated by sequencing, and then mapped (Map_with_Bowtie_for_Illumina_V1.1.2) to the *C. crescentus* NA1000 genome (NC_011916.1) using the web-based analysis platform Galaxy (https://usegalaxy.org). Using Samtool_V0.1.18, BED file format encompassing the Tn insertion coordinates were generated and then imported into SeqMonk V1.40.0 (www.bioinformatics.babraham.ac.uk/projects/) to assess the total number of Tn insertion per chromosome position (Tn-insertion per millions of reads count) or per coding sequence (CDS). For CDS Tn-insertion ratio calculation, SeqMonk datasets were exported into Microsoft Excel files (Dataset Table S6) for further analyses and used to generate Figure S4B, as described previously^22^. Briefly, to circumvent ratio issue for a CDS Tn-insertion value of 0 and CDS that do not share sufficient statistical Tn-insertions, an average value of all CDS-Tn insertions normalized to the gene size was calculated, and 1% of this normalized value was used to correct each CDS-Tn insertion value.

#### NA1000, NA1000 ctrA401 and ctrA(T170A) Tn-Seq

An overnight *WT* (NA1000), *ctrA401* (T170I) and *ctrA(T170A)* cell cultures were grown in PYE and mutagenized with a *himar1* transposon (Tn)^23^. Transposon mutagenesis of *C. crescentus* NA1000, NA1000 *ctrA401* and *ctrA(T170A)* was done by mobilizing the *himar1* transposon (KanR) from plasmid pHPV414 in *Escherichia coli* S17-1 λ*pir* into the *WT* (NA1000), *ctrA401* and *ctrA(T170A)* strain and selecting for kanamycin-nalidixic acid-resistant *C. crescentus* clones. For the *C. crescentus* NA1000, NA1000 *ctrA401*::Tn(Kan^R^) and *ctrA(T170A)*::Tn(Kan^R^), Tn-libraries of >200,000 kanamycin-resistant clones were collected. *C. crescentus* NA1000, NA1000 *ctrA401*::Tn and *ctrA(T170A)*::Tn banks were grown overnight in PYE and chromosomal DNA was extracted. Genomic DNA was used to generate barcoded Tn-Seq libraries and submitted to Illumina HiSeq 2000 sequencing (Fasteris SA). Tn-insertion specific reads (50 bp long) were sequenced using the himar-Tnseq1 primer (5’-AGACCGGGGACTTATCAGCCAACCTGT-3’) and yielded several million reads that were mapped to *Caulobacter crescentus* NA1000 (NC_011916.1) as previously describes^22^. CDS Tn-insertions calculation are presented in Table S7 and were used to generate Figure 3E and S6.

### Stalk suppressors of the *ctrA401* motility mutant cells

To isolate mutation that restores the polarity of the *ctrA401* MS2 mutant cells, we integrated at the chromosomal *pilA* locus the transcriptional reporter P*_pilA_-nptII* and select for suppressors that grow on PYE agar plates supplemented with 40 μg/mL of kanamycin. Kanamycin resistant clones were then screened by DIC microscopy for the presence of stalk. Five isolates restored the stalk synthesis and were subjected to Sanger sequencing for the *ctrA* gene to exclude potential revertant. Sanger sequencing reveals instead of revertant, a second mutation within *ctrA* in all the five isolates, the *ctrA(T168I)*, in which the codon of threonine (ACC) at residue 168 was exchanged for one encoding an isoleucine (ATC). To confirm that this mutation was responsible for the overhaul of stalk synthesis, we backcross the both mutations (T170I and T168I) into the *WT* cells. To this end, a pNPTS138-*ctrA*-ds, conferring kanamycin resistance, was integrated by homologous recombination downstream of the *ctrA* gene in the *ctrA401* MS2 *pilA*::P*_pilA_*-*nptII* SS mutant. Next, ϕCr30-mediated generalized transduction was used to transfer the *ctrA(T168I/T170I)* allele into the *WT* cells. Sucrose 3% was then used to eliminate the pNPTS138-*ctrA*-ds by homologous recombination and clones were screen for kanamycin susceptibility. Sanger sequencing was used to verify the integrity of the mutation using primers ctrA_NdeI and ctrA_XbaI and the resulting mutants were probed for the presence of the stalk by phase contrast microscopy.

### β-Galactosidase assays

β-Galactosidase assays were performed at 30°C. Cells (50 to 200 µL) at OD_660nm_ =0.1-0.5 were lysed with chloroform and mixed with Z buffer (60 mM Na_2_HPO_4_, 40 mM NaH_2_PO_4_, 10 mM KCl and 1 mM MgSO_4_ heptahydrate) to a final volume 800 µL. Two hundred µL of ONPG (4 mg/mL o-nitrophenyl-β-D-galactopyranoside in 0.1 M KPO_4_ pH7.0) was added and the reaction timed. When a medium-yellow colour developed, the reaction was stopped with 400 µL of 1M Na_2_CO_3_. The OD_420nm_ of the supernatant was determined and the units were calculated with the equation: U= (OD_420nm_ * 1000) / (OD_660nm_ * time (in min) * volume of culture (in mL)). Experimental values represent the averages of 4 independent experiments and error was computed as standard deviation (SD).

### Immunoblot analysis

Protein samples were separated by SDS-PAGE and blotted on PVDF (polyvinylidenfluoride) membranes (Merck Millipore). Membranes were blocked for 1h with TBS, 0,1% Tween 20 and 5% dry milk and then incubated for overnight with the primary antibodies diluted in TBS, 0,1% Tween 20, 5% dry milk. The different antisera were used at the following dilutions: anti-SciP (1:5000)^15^, anti-FljK (1:50000)^24^, anti-CtrA (1:20000) (Ardissone, unpublished), anti-PilA (1:10000)^25^, anti-HfsJ (1:10000) (Eroglu, unpublished), anti-SpmX (1:50000)^26^ and anti-CCNA_00163 (1:10000)^5^ or anti-MreB (1:200000)^27^ as a loading control. The membranes were washed 4 times for 5 min in TBS, 0.1% Tween 20 and incubated for 1h with the secondary antibody diluted in TBS, 0.1% Tween 20 and 5% dry milk. The membranes were finally washed again 4 times for 5 min in in TBS, 0.1% Tween 20 and revealed with Immobilon Western Blotting Chemoluminescence HRP substrate (Merck Millipore) and Super RX-film (Fujifilm).

### Microscopy

PYE cultivated cells in exponential or stationary growth phase were immobilized using a thin layer of 1% agarose. DIC and Phase microscopy images were taken with an Alpha Plan-Apochromatic 100X/1.46 DIC(UV) VIS-IR oil and Alpha Plan-Apochromatic 100X/1.46 Ph3(UV) VIS-IR oil objectives respectively on an Axio Imager M2 microscope (Zeiss) with a 405 and 488 nm lasers (Visitron Systems GmbH, Puchheim, Germany) and a Photometrics Evolve camera (Photometrics) controlled through Metamorph V7.5 (Universal Imaging). Images were processed using Metamorph V7.5 and ImageJ.

### Flow cytometry analysis

PYE cultivated cells in exponential (OD_660nm_ =0.3-0.6) and stationary growth phase were fixed into ice cold 77% Ethanol solution. Fixed cells were resuspended in FACS Staining buffer pH7.2 (10 mM Tris-HCl, 1 mM EDTA, 50 mM NaCitrate, 0.01% TritonX-100) and then treated with RNase A (Roche) at 0.1 mg/mL during 30 min at room temperature. Cells were stained in FACS Staining buffer containing 0.5 µM of SYTOX Green nucleic acid stain solution (Invitrogen) and then analysed using a BD Accuri C6 flow cytometer instrument (BD Biosciences, San Jose, California, USA). Flow cytometry data were acquired using the CFlow Plus V1.0.264.15 software (Accuri Cytometers Inc.) and analysed using FlowJO software. 20000 cells were analysed from each biological sample. The Green fluorescence (FL1-A) parameters were used to estimate cell chromosome contents. Relative chromosome number was directly estimated from FL1-A value of *WT* cells treated with 30 µg/mL Rifampicin during 3 hours at 30°C. Rifampicin treatment of cells blocks the initiation of chromosomal replication but allows ongoing rounds of replication to finish. Each experiment was repeated independently, and representative results are shown.

### Motility assays and phage infectivity tests

Swarming properties were assessed with 1.5 μL drops of overnight culture, adjusted to an OD_660nm_ of 1, spotted on PYE soft agar plates (0.3% agar) and grown for 48 hours. Phage susceptibility assays were conducted by mixing 300 μL of overnight culture in 6 mL soft PYE agar and overlaid on a PYE agar plate. Upon solidification of the soft (top) agar, 4 μL drops of serial dilution of phages (ϕCbK or ϕCr30) were spotted and scored for plaques after one day incubation at 4°C.

### Viability test on plates

Viability of *WT*, *ctrA401* and derivatives was assessed with 4 μL drops of serial dilutions of liquid culture (10^−1^ to 10^−6^), adjusted to OD_660nm_=1, on PYE plates, then incubated for 2 days at 30°C or 37°C.

### Chromatin ImmunoPrecipitation coupled to deep Sequencing (ChIP-Seq)

Mid-log phase and stationary phase cells cultured in PYE were cross-linked in 10mM sodium phosphate (pH 7.6) and 1% formaldehyde at room temperature for 10 min and on ice for 30 min thereafter and washed three times in phosphate-buffered saline (PBS). Cells were lysed in a Ready-Lyse lysozyme solution (Epicentre Technologies) according to manufacturer’s instructions and lysates were sonicated in an ice-water bath for 15 cycles 30 s ON and 30 s OFF, to shear DNA fragments to an average length of 0.3-0.5 kbp and cleared by centrifugation at 14000 g for 2 min at 4°C. Lysates were normalized by protein content, diluted to 1 ml using ChIP buffer (0.01% SDS, 1.1% Triton X-100, 1.2 mM EDTA, 16.7 mM Tris-HCl (pH 8.1), 167 mM NaCl plus protease inhibitors (Roche, Switzerland) and pre-cleared with 80 μL of protein-A (rabbit antibodies) agarose (Roche) and 100 μg BSA. Ten percent of the supernatant was removed and used as total chromatin input DNA as described before^26^.

Two microliters of rabbit polyclonal antibodies to CtrA (Ardissone) were added to the remains of the supernatant, incubated overnight at 4 °C with 80 μL of protein-A beads pre-saturated with BSA, washed once with low salt buffer (0.1% SDS, 1% Triton X-100, 2 mM EDTA, 20 mM Tris-HCl (pH 8.1) and 150 mM NaCl), high salt buffer (0.1% SDS, 1% Triton X-100, 2 mM EDTA, 20 mM Tris-HCl (pH 8.1) and 500 mM NaCl) and LiCl buffer (0.25 M LiCl, 1% NP-40, 1% sodium deoxycholate, 1 mM EDTA and 10 mM Tris-HCl (pH 8.1)), and twice with TE buffer (10 mM Tris-HCl (pH 8.1) and 1 mM EDTA). The protein DNA complexes were eluted in 500 μL freshly prepared elution buffer (1% SDS and 0.1 M NaHCO3), supplemented with NaCl to a final concentration of 300 mM and incubated overnight at 65 °C to reverse the crosslinks. The samples were treated with 2 μg of Proteinase K for 2 h at 45 °C in 40 mM EDTA and 40 mM Tris-HCl (pH 6.5). DNA was extracted using phenol:chloroform:isoamyl alcohol (25:24:1), ethanol precipitated using 20 μg of glycogen as carrier and resuspended in 50 μL of water.

Immunoprecipitated chromatin was used to prepare sample libraries used for deep-sequencing at Fasteris SA (Geneva, Switzerland). ChIP-Seq libraries were prepared using the DNA Sample Prep Kit (Illumina) following the manufacturer’s instructions. Single-end run were performed on a Next-Generation DNA sequencing instrument (NGS, HiSeq3000/4000), 50 cycles were read and yielded several million reads. The single-end sequence reads stored in FastQ files were mapped against the genome of *C. crescentus* NA1000 (NC_011916.1) and converted to SAM using BWA and SAM tools respectively from the galaxy server (https://usegalaxy.org/). The resulting SAM was imported into SeqMonk (http://www.bioinformatics.babraham.ac.uk/projects/seqmonk/,version1.45.1) to build sequence read profiles. The initial quantification of the sequencing data was done in SeqMonk: the genome was subdivided into 50 bp probes, and for every probe we calculated a value that represents a normalized read number per million (Supplemental tables S2, S3 and S4).

Using the web-based analysis platform Galaxy (https://usegalaxy.org), CtrA ChIP-Seq peaks were called using MACS2^28^ relative to the total input DNA samples. The q-value (false discovery rate, FDR) cut-off for called peaks was 0.05. Peaks were rank-ordered according to fold-enrichment (Suppl. Table S5) and peaks with a fold-enrichment values >2 were retained for further analysis.

The heatmaps in Figure S1 were generated using SeqMonk and ChIP-seq signals were plotted over 800 bp ranges at each *C. crescentus* Coding DNA Sequence (CDS) centered at each translation initiation codon. Sequence data have been deposited to the Gene Expression Omnibus (GEO) database.

### Molecular modelling of CtrA

Models of CtrA, CtrA(T170I) and CtrA(T168I/T170I) homo-dimers were generated based on the crystal structure of the DNA-binding transcriptional regulator BasR/PmrA from *Klebsiella pneumoniae* (PDB 4s05.1.B), using the SWISS-MODEL server^29^. Comparison with the solved structured of *Brucella abortus* receiver domain (PDB 4QPJ^30^) was performed with the built model to control the quality of the model. Pymol was used to visualize and map the location of the mutated amino acids within the structure.

## Results

### CtrA401 uncouples the switch from late-S-phase to G1-phase promoters

We confirmed that the CtrA-activated promoter P*_fliQ_* is upregulated in *ctrA401* cells at 30°C^6^ using a P*_fliQ_*-*lacZ* promoter probe plasmid in which P*_fliQ_* is transcriptionally fused to the promoterless *lacZ* gene. We observed a more than two-fold increase in LacZ activity in *ctrA401* cells (241%, Table 2) versus *WT* cells growing exponentially at 30°C. To determine if other CtrA-activated promoters are upregulated in the same manner, we conducted LacZ-based promoter probe assays in *WT* and *ctrA401* cells at 30°C (Table 2). Remarkably, we observed a similar behavior with promoters that CtrA activates in late S-phase (e.g. P*_fliX,_* P*_fliL,_* P*_pleA_*, P*_cpaB_* and P*_CCNA_03790_*; Table 2), but not with promoters that CtrA activates in G1-phase (e.g. P*_pilA_*, P*_flaF,_* P*_hfsJ,_* P*_CCNA_02061_* or P*_sciP_*; Table 2). In fact, promoters of the G1 class were poorly active in the *ctrA401* background and immunoblotting revealed that proteins expressed from G1 phase promoters that depend directly (HfsJ, PilA, SciP; Figure 1B, S1B, S1D and S2B) or indirectly (SpmX; Figure 1B) on CtrA are not (or poorly) expressed in *ctrA401* cells. These results suggest that *ctrA401* cells are unable to activate these G1-phase promoters, but efficiently induce the late S-phase promoter class.

**Table 2.** Activity of CtrA-activated promoters in C. crescentus WT & mutants. Promoter-probe assays of transcriptional reporters carrying a various *lacZ-* based CtrA dependent promoter in *WT* and *ctrA401* cells and derivatives. Black refers to late S-phase promoters and blue to the G1-phase promoters. *spmX* is represented in purple as it is not a direct target of CtrA. Values are expressed as percentages (activity in *WT* set at 100%). Data from four independent experiments, error bars are standard deviation. ND, Not determined.

| Strain : | WT | $\Delta spoT$ | *ctrA401* | *ctrA401*<br>RT | $\Delta spoT$<br>*ctrA401* | *ctrA401*<br>MS2 | *ctrA401*<br>MS4 | *ctrA401*<br>HS |
| --- | --- | --- | --- | --- | --- | --- | --- | --- |
| Promoter : |  |  |  |  |  |  |  |  |
| *fliL* | 100 ± 8 | 115 ± 3 | 175 ± 4 | 178 ± 2 | 145 ± 9 | 161 ± 5 | 153 ± 9 | 145 ± 4 |
| *fliQ* | 100 ± 3 | 87 ± 2 | 241 ± 2 | 237 ± 2 | 201 ± 4 | 249 ± 6 | 218 ± 6 | 137 ± 2 |
| *fliX* | 100 ± 2 | 93 ± 3 | 131 ± 6 | 128 ± 3 | 110 ± 5 | 124 ± 7 | 129 ± 6 | 128 ± 3 |
| *pleA* | 100 ± 5 | 93 ± 4 | 235 ± 9 | 239 ± 12 | 199 ± 20 | 231 ± 11 | 216 ± 16 | 107 ± 8 |
| **CCNA_03790** | 100 ± 6 | 86 ± 5 | 131 ± 10 | 133 ± 8 | 134 ± 10 | 134 ± 7 | 129 ± 10 | 95 ± 6 |
| *ctrA* | 100 ± 3 | 84 ± 1 | 90 ± 4 | 88 ± 2 | 78 ± 2 | 87 ± 3 | 90 ± 2 | 127 ± 1 |
| *ccrM* | 100 ± 3 | 85 ± 1 | 70 ± 2 | 73 ± 1 | 64 ± 1 | 62 ± 2 | 60 ± 1 | 78 ± 5 |
| *sciP* | 100 ± 2 | 95 ± 4 | 30 ± 3 | 28 ± 3 | 36 ± 2 | 25 ± 3 | 28 ± 3 | 58 ± 3 |
| *hfsJ* | 100 ± 2 | 85 ± 1 | 19 ± 1 | 21 ± 2 | 21 ± 1 | 19 ± 1 | 18 ± 1 | 54 ± 3 |
| **CCNA_02061** | 100 ± 8 | 82 ± 10 | 29 ± 5 | 29 ± 4 | 29 ± 4 | 32 ± 5 | 36 ± 5 | 55 ± 2 |
| *flaF* | 100 ± 2 | 77 ± 3 | 66 ± 1 | 71 ± 2 | 72 ± 2 | 70 ± 3 | 60 ± 2 | 75 ± 3 |
| *pilA* | 100 ± 1 | 95 ± 7 | 20 ± 2 | 21 ± 1 | ND | 16 ± 1 | 17 ± 1 | 59 ± 1 |
| *spmX* | 100 ± 2 | 102 ± 2 | 36 ± 1 | 29 ± 1 | 33 ± 3 | 27 ± 1 | 29 ± 3 | 91 ± 3 |

To probe for commensurate changes in promoter occupancy of CtrA401 (T170I) versus WT CtrA, we conducted <u>ch</u>romatin-<u>i</u>mmuno<u>p</u>recipitation followed by deep-<u>seq</u>uencing (ChIP-Seq; Figure 1C, S1A) analyses of CtrA-bound sites. Immunoprecipitations were done by incubating polyclonal antibodies to CtrA with chromatin extracted from *WT* and *ctrA401* cells (grown exponentially at 30°C). Deep-sequencing revealed that the chromosomal occupancy of CtrA401 (T170I) is generally elevated compared to that of CtrA (WT), without noticeably affecting CtrA abundance inside cells (Figure 1C, inset). Thus, the T170I mutation does not compromise general binding to the CtrA box *in vivo*. As expected, the specific association of CtrA401 with late S-phase promoters such as P*_fliQ_*, P*_cpaB_* and P*_fliF_* is increased compared to that of CtrA (WT; Figure 1D), while at G1-phase promoters, such as P*_sciP_*, P*_hvyA_*, P*_hfsJ_*, P*_tacA_* and P*_pilA_* (Figure 1D), CtrA occupancy is clearly diminished. Moreover, a control promoter that CtrA does not directly target (the promoter of the *fljL* flagellin gene) showed poor occupancy of CtrA *in vivo* and little or no difference between *WT* and *ctrA401* cells (Figure 1D, note change in scale).

We conclude that CtrA401 is impaired in the switch from late-S-phase to G1 phase promoter activation and binding. Since this defect also results in a failure to express the negative regulator of late S-phase promoters^12, 15, 16^, SciP (Figure 4D, 4E, S1C, S1D), these promoters are no longer shut off and firing is maintained, resulting in elevated activity. Thus, the negative regulation of CtrA on late S-phase target promoters such as P*_fliQ_* is indirect, stemming from a defect in activating SciP expression in *ctrA401* cells which explains why this allele of *ctrA* surfaced in the previous selection for loss of negative regulation at P*_fliQ_*^6^. Importantly, and contrary to earlier interpretations of the phenotype of *ctrA401* cells, our findings suggest that the phenotypes observed at the permissive temperature are not due to a general loss of CtrA function/binding at target promoters, but is instead due to a specific defect due to crippled G1-phase promoter firing and the resulting overactivation of late S-phase promoters.

### Capsule inhibits swarming motility of *ctrA401* cells

Since the G1-phase is the principal phase of motility in *C. crescentus*, motility assays serve as proxy for perturbations in G1-phase duration (and gene expression). Consistent with the defect in G1-phase promoter activation by the *ctrA401* mutation, these cells are poorly motile on swarm (0.3%) agar plates (Figure 2B) even though early flagellar promoter fire at an elevated level (Table 2; Figure 2A). Moreover, immunoblotting revealed that the FljK flagellin, a late flagellar component that is assembled into the flagellar filament, is also more abundant in *ctrA401* cells than in *WT* cells (Figure 2B, S2A). Since a block in flagellar assembly prevents expression of FljK^31^, these immunoblotting experiments indicate that *ctrA401* cells do not suffer from a flagellar assembly block and, thus, that swarming motility is impaired in *ctrA401* cells for (an)other reason(s).

**Figure 2.**
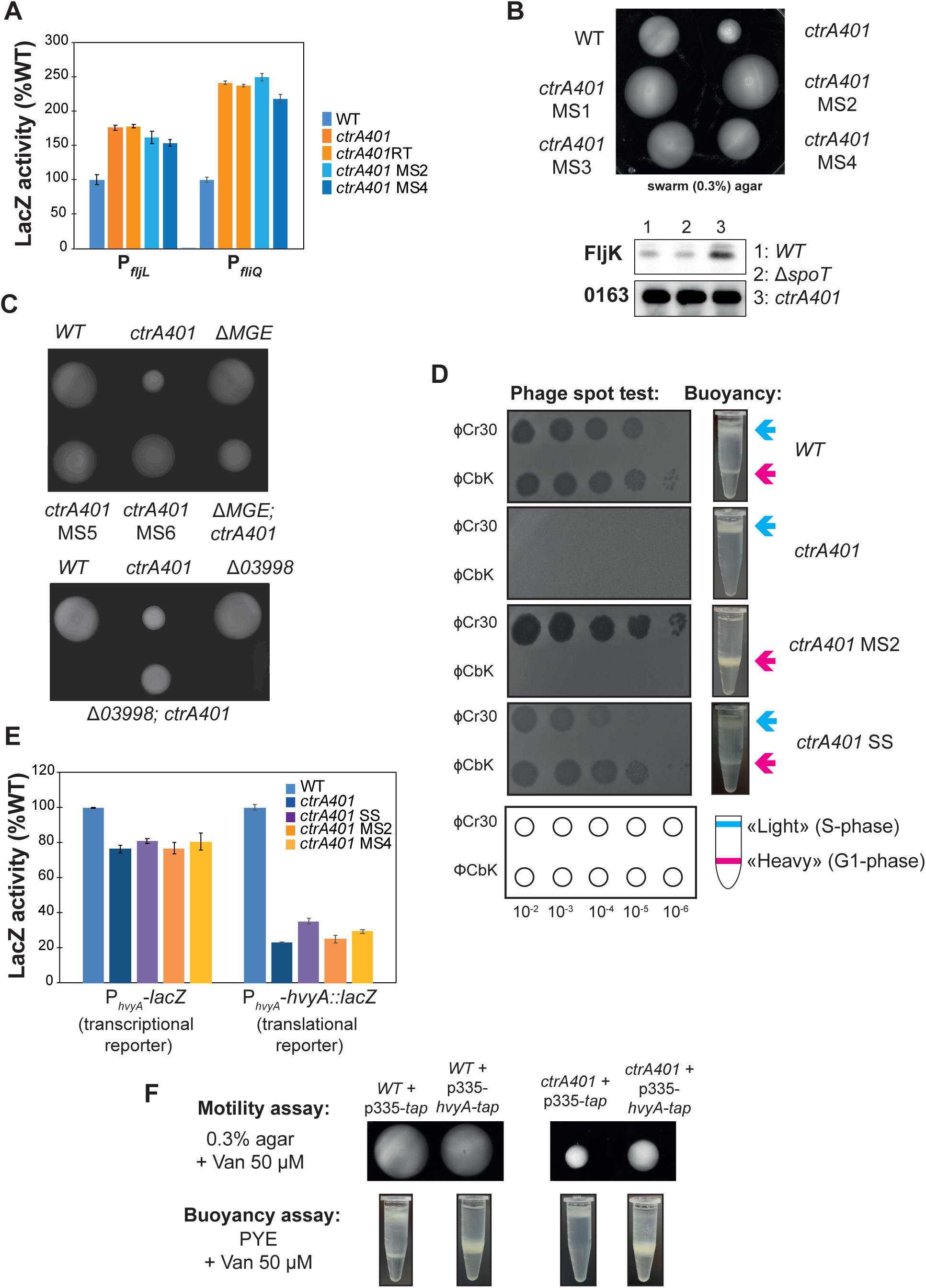
Capsulation of the *ctrA401* mutant affects buoyancy switch, bacteriophage sensitivity and motility. A) Promoter-probe assays of transcriptional reporters carrying a *fljL*, *fliQ* (class IV and class II genes, respectively) promoters in *WT*, *ctrA401* and derivatives. Values are expressed as percentages (activity in *WT* set at 100%). B) Motility plates (0,3% agar) inoculated with *WT*, *ctrA401*, *ctrA401* motility suppressors 1 and 2 (*ctrA401-MS1* and *ctrA401-MS2*) strains and immunoblot showing the steady state levels of the alpha-flagellin FljK in *WT*, Δ*spoT* and *ctrA401* cells. C) Swarm (0.3%) agar inoculated with *WT*, *ctrA401*, *ctrA401-MS*, Δ*MGE* and Δ*03998*. D) Schematic and pictures of cell buoyancy upon centrifugation on density gradient and sensitivity to bacteriophages ϕCr30 and ϕCbK for *WT* Caulobacter and *ctrA401* mutants and derivatives. E) Promoter-probe assays of *hvyA*-transcriptional (left graph) and translational reporters (right graph) in *WT*, *ctrA401* and derivatives. Values are expressed as percentages (activity in *WT* set at 100%). Data from four independent experiments, error bars are standard deviation. F) Swarming motility and buoyancy assays in cells overexpressing *hvyA-tap* fusion under control of P*van* on *pMT335* plasmid restores the “heavy” cell buoyancy to the *ctrA401* cells.

To determine the basis of this motility defect, we isolated 10 <u>m</u>otile <u>s</u>uppressors (*ctrA401-MS*) emanating as flares on swarm agar from a background inoculum of (non-motile) *ctrA401* cells (Figure 2B, 2C). Promoter probe assays with several early (class II) flagellar and other promoters in *ctrA401-MS*2 and *ctrA401-MS*4 cells (Table 2, Figure 2A, S3A) showed that CtrA-dependent regulation is not substantially altered in these mutants compared to *ctrA401* parental cells. Thus, the suppressor mutation in *ctrA401-MS* cells lies outside of the CtrA regulatory hierarchy. Surprisingly, *ctrA401-MS* cells rapidly sediment to the bottom of the cultivation tube, whereas *ctrA401* parental or *WT* cultures remained in suspension during the same interval (Figure 2D). This sedimentation phenotype resembles that of capsule-less mutants that have a reduced cellular buoyancy^5^. Indeed, buoyancy analyses by Percoll density gradient centrifugation revealed that *ctrA401-MS*2 cultures only contained “heavy” cells (Figure 2D), whereas *ctrA401* cells were only “light”. As *WT* cultures contain both “heavy” (non-capsulated) G1-phase cells and “light” (capsulated) S-phase cells (Figure 2D), we reasoned that *ctrA401* cells remain “light” because of a failure to activate a G1-phase promoter and that this block also perturbs swarming motility.

The CtrA-activated promoter of the *hvyA* gene, encoding a G1-specific capsulation inhibitor, fires in G1-phase (P*_hvyA_*)^12^. Loss of HvyA leads to constitutive capsulation and thus only “light” cells in the culture^5^. After confirming that a transcriptional fusion to the P*_hvyA_*-*lacZ* and a P*_hvyA_*-*hvyA*::*lacZ* translational fusion is poorly active in *ctrA401* cells (Figure 2E), we reasoned that the constitutive capsulation phenotype of *ctrA401* cells is due to a failure in expressing HvyA. Indeed, constitutive expression of an epitope-tagged version of HvyA bearing a C-terminal <u>T</u>andem <u>A</u>ffinity <u>P</u>urification (TAP) from a vanillate-inducible promoter on the pBBR-based plasmid pMT335^32^ not only reversed the buoyancy defect of *ctrA401* cells (Figure 2F), but also improved swarming motility.

If *ctrA401* cells are always capsulated and “light”, then the “heavy” *ctrA401 MS* cells should lack capsule. Since capsule protects cells from infection by ϕCr30 by masking its receptor, the S-layer protein RsaA^33^, we predicted that *ctrA401* cells should be resistant to infection by ϕCr30, whereas *ctrA401-MS*2 should be hypersensitive to ϕCr30. The phage spot (lysis) tests shown in Figure 2D confirmed this prediction. As control for specificity we also conducted phage spot tests with caulophage ϕCbK which requires pili for infection^34^. Both *ctrA401* and *ctrA401-MS*2 mutants are resistant to ϕCbK because they do not express the *pilA* pilin gene encoding the structural subunit of the pilus filament that is expressed from the CtrA activated P*_pilA_* promoter in G1-phase (Table 2, Figure S1B, S2B). By contrast, *WT* cells are piliated and thus infected (and lysed) by ϕCbK.

Why might the *ctrA401-MS* mutants have lost the ability to form capsule? We suspected the acquisition of a mutation in a capsule biosynthesis gene in these mutants. A cluster of capsule biosynthetic proteins is encoded on a <u>m</u>obile <u>g</u>enetic <u>e</u>lement (MGE; Figure S3B) cells that can sporadically excise^5, 35^. PCR-based analyses of the *ctrA401*-*MS* mutants revealed that the MGE had been lost from *ctrA401-MS*4-7 cells as indicated by the increased abundance of the *attB* PCR product that is amplified when the *attB* junction is created by MGE excision (Figure S3B, S3C), whereas the MGE is present in *WT* (NA1000) and *ctrA401* cells. Further analysis by PCR revealed that *ctrA401-MS*4-7 cells lacked the capsule genes (*CCNA_03998*) and the *CCNA_00477* encoded on the MGE. In the remaining mutants, *CCNA_0477* encoded on the MGE is present, but *CCNA_03998* was not detected in *ctrA401-MS*2 and a larger rearrangement of the *CCNA_03998* had occurred in *ctrA401-MS*1 (Figure S3C). Prompted by the observation that *CCNA_03998* is rearranged or absent in the aforementioned *ctrA401*-*MS* mutants, we asked if the remaining four MS mutants (*MS3*, *8*, *9*, *10*) had acquired point mutations in *CCNA_03998* that would impair function. Indeed, PCR-sequencing revealed that *ctrA401-MS*3 has a nonsense mutation at codon 61 (changing the tryptophan codon to a stop codon), while *ctrA401-MS*8 and *ctrA401-MS*9 have an identical mutation at codon 331, changing a cysteine codon into a stop codon. The remaining mutant, *ctrA401-MS*10, has a missense mutation at codon 32, changing the proline codon to a histidine codon (Figure S3D).

Taken together these analyses suggest that loss of the MGE and, specifically, loss of capsule biosynthetic function of the CCNA_03998 WbuB-like glycosyltransferase, promotes swarming motility of *ctrA401* cells. In support of the conclusion that constitutive capsulation interferes with swarming motility and likely flagellar function in *ctrA401* cells, we transduced the *ctrA401* mutation into ΔMGE cells as well as Δ*CCNA_03998* mutant cells and found that swarming motility is indeed improved in the resulting double mutants compared to the *ctrA401* parent (Figure S3E). However, motility is reduced in these ΔMGE; *ctrA401* mutants compared to *ctrA401-MS* mutants, presumably because of yet unknown mechanism of motility inhibition in *ctrA401* or because of unknown contributions conferred by the suppressor mutations on the MGE.

### Intragenic suppressor mutations of *ctrA401* alter DNA-binding

To investigate the G1-promoter block of *ctrA401* cells further, we designed another suppressor screen to select for ameliorated G1-phase gene expression using the *pilA*::P*_pilA_*-*nptII* transcriptional reporter conferring resistance to kanamycin^12, 36^. For practical reasons, we chose *ctrA401-MS*2 as host for the reporter since this mutant showed the same regulatory defects on CtrA-dependent gene expression as *ctrA401* parent cells (Table 2), with the added benefit of enabling convenient transfer of the *pilA*::P*_pilA_*-*nptII* reporter using ϕCr30-mediated generalized transduction, which is not possible for *ctrA401* parental cells (see above). The *pilA*::P*_pilA_*-*nptII* reporter confers resistance to 40 μg/mL kanamycin (on plates) to *WT* cells (Figure S4A). As expected, *pilA*::P*_pilA_*-*nptII* is poorly active in *ctrA401-MS*2 cells, precluding growth on kanamycin (40 μg/mL) plates (Figure S4A). In an attempt to identify extragenic suppressor mutations that restore G1-phase promoter activity to *ctrA401* cells, we conducted transposon (Tn) mutagenesis of the reporter strain using a *himar1* Tn conferring resistance to gentamicin. Selecting for resistance to gentamicin and kanamycin of *ctrA401*-*MS*2 *pilA*::P*_pilA_*-*nptII* reporter cells following Tn mutagenesis, revealed several clones with Tn insertions at different positions within or near the *pilA* promoter (Figure S4B, S4C, Table S6), suggesting that outward transcription form the Tn bypasses the dependency for CtrA to activate P*_pilA_*-*nptII*.

As mapping of over 20 different Tn mutants failed to reveal Tn insertions in *loci* other than P*_pilA_*, we sought intragenic suppressor mutations in *ctrA401* that allow growth of *ctrA401*-*MS*2 *pilA*::P*_pilA_*-*nptII* reporter cells on plates containing kanamycin (40 μg/mL). To eliminate isolating mutants harboring promoter-up mutations in P*_pilA_* that permit growth on kanamycin plates but that do not otherwise impact the CtrA regulon, we imposed a second visual screen for stalks in the kanamycin-resistant mutants, since the polar stalk is absent from *ctrA401* cells^6^ and it can be conveniently recognized by differential interference contrast (DIC) microscopy. As shown in Figure 3A, *ctrA401* and *ctrA401*-*MS*2 *pilA*::P*_pilA_*-*nptII* parental cells are stalkless when grown in PYE, but stalks are produced in the *ctrA401-SS* (<u>s</u>talked <u>s</u>uppressor) mutant. Moreover, DIC microscopy and fluorescence-activated cell sorting (FACS) revealed that the cell filamentation defect is also ameliorated in the SS mutant cells compared to *ctrA401* and *ctrA401*-*MS*2 *pilA*::P*_pilA_*-*nptII* parental cells (Figure 3A).

**Figure 3.**
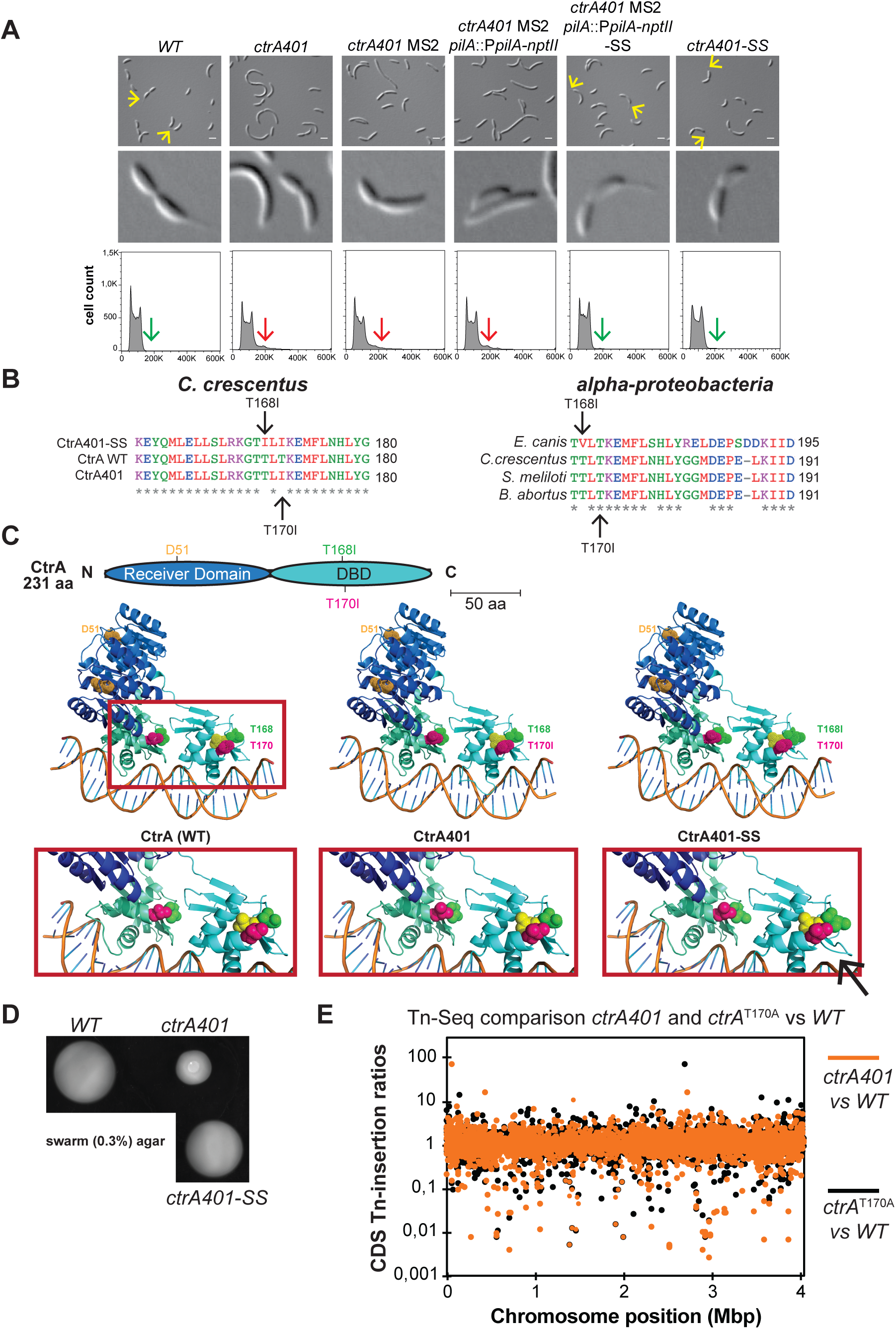
Intragenic suppressor mutations of *ctrA401* alter DNA-binding. A) DIC microscopy, 8X zoom images and FACS analysis of the *WT, ctrA401* and derivatives carrying the *pilA::PpilA-nptII* transcriptional reporter. Microscopy pictures and DNA content (FL1-A channel) quantification were performed during exponential growth in PYE. *ctrA401 MS2 pilA::PpilA-nptII* cells resistant to 40 µg/mL of kanamycin (kan) were screened by microscopy for the presence of stalk (yellow arrow). DIC images and FACS analysis of the *ctrA401* and *ctrA401-MS2* show cells that are elongated, accumulating more than 2n chromosomes (red arrow) and stalkless in exponential phase. Selection on PYE solid supplemented with 40 µg/mL of kanamycin of *ctrA401 MS2 pilA::PpilA-nptII* resistant cells and screening by DIC microscopy for the presence of stalk (yellow arrow, 8X zoom) lead to the identification of the *ctrA401-SS* (T170I/T168I) mutation. *ctrA401-SS* cells do not elongate and do not accumulate extra-chromosome (green arrow). B) On the left, the alignment of the 28 amino acids around the T170 and the T168 shows the amino acids replacement in *ctrA401*, *ctrA401-SS* compared to *WT*. On the right, the alignment of CtrA protein sequences of several members of the alpha-proteobacteria phylum is shown. C) Schematic of CtrA domains and models of CtrA, CtrA401 and CtrA401-SS homodimers structures based on homology together with promoter DNA. Receiver domains are indicated in dark blue and DNA-binding domains (DBD) are indicated in light blue. D51, that constitutes the phosphorylation site of CtrA, is indicated as sphere in orange, the T168 (CtrA, CtrA401) and the I168 (CtrA401-SS) are indicated as sphere in green and the T170 (CtrA) or the I170 (CtrA401, CtrA401-SS) as sphere in pink. L169 is shown as sphere in yellow. Red rectangle shows a zoom within the structure and the proximity of T168 to the ß7- ß8 of the second DBD and the change observed in the C-terminal ß-hairpin (black arrows) in CtrA-SS. D) Motility plates (0,3% agar) inoculated with *WT*, *ctrA401*, *ctrA401-SS* cells. The T168I amino acid change helps the *ctrA401* to restore motility. E) Tn insertion bias in coding sequences (CDS) of *ctrA401* (T170I) cells relative to *WT* cells (orange dots) and *ctrA*(T170A) cells relative to WT (black dots) cells as determined by Tn-seq. Abscissa shows position as function of genome position, and ordinate gives Tn-insertion ratio. Peaks show CDSs with the highest number of Tn insertions. Noncoding sequences are not included.

Backcrossing experiments and DNA sequencing revealed that the *ctrA401-SS* mutant had acquired an additional mutation in *ctrA*, changing the threonine codon 168 into an isoleucine codon (T168I). T168I lies two residues ahead of the T170I mutation in CtrA401 (Figure 3B) within the DNA-binding domain of CtrA. The stretch from 167-191 in *C. crescentus* CtrA is highly conserved in the alpha-proteobacteria (Figure 3B), but strikingly T168 is also substituted by a hydrophobic residue (valine) at the corresponding position in the rickettsial lineage (*Ehrlichia canis* in Figure 3B), suggesting that this position in CtrA likely reflects a key residue in determining the DNA-binding activity of CtrA at G1-phase promoters in free-living alpha-proteobacteria and developmental (growth phase transition) control in the obligate intracellular lineage.

Modelling the promoter bound structure of WT and mutant CtrA (Figure 3C), along with ChIP-Seq analysis of *ctrA401-SS* cells (obtained from backcrossing the *ctrA401-SS* allele into *WT* cells, Figure 1C) supported the key role of this residue. The CtrA401-SS (T168I/T170I) protein shows an increase in occupancy at G1-phase promoters and a reduction at late S-phase promoters, thus correcting the properties of the CtrA401 (T170I) mutant towards the *WT* properties (Figure 1C, 1D). Importantly, promoter probe assays confirmed that promoter firing was also ameliorated in the *ctrA401-SS* mutant compared to *ctrA401* cells: while firing is G1-phase elevated compared to the *ctrA401* parent (Table 2), expectedly late S-phase promoters are down-regulated since P*_sciP_* activity is also elevated *ctrA401-SS* compared *ctrA401* cells. Immunoblotting confirmed that the steady-state levels of HfsJ and SpmX both directly (HfsJ) or indirectly (SpmX) dependent on G1-phase promoters activated by CtrA, are restored to near WT levels in *ctrA401-SS* cells compared to *ctrA401* cells (Figure 1B).

In addition to stalk formation, the other misregulated phenotypes of *ctrA401* cells are also ameliorated by the *ctrA401-SS* mutation, including the resistance to ϕCbK (i.e. piliation, Figure 2D, S1B), as well as the resistance to ϕCr30 along with the exclusive “light” buoyancy (i.e. constitutive capsulation, Figure 2D) phenotype. As expected, since the *ctrA401-SS* mutation allows the formation of non-capsulated G1 cells because of the improved expression of HvyA, motility is also improved compared to *ctrA401* parental cells (Figure 3D). Importantly, *ctrA401-SS* cells have an elevated heat tolerance (plating efficiency at 37°C) compared to *ctrA401* cells (Figure S1C).

Thus, residues T170 and T168 define the G1 promoter binding capacity of CtrA at G1-phase promoters and confer opposing phenotypes depending on the side chain replacement. This notion is further reinforced by the previously finding of suppressor mutations in these residues (i.e. T168A and T170A) with the opposite effect, restoring motility to cells in which G1-promoters are deprepressed^12^. Determining the transposon (Tn) insertions profile by Tn-Seq (transposon insertion sequencing) analysis of *ctrA401* (T170I) and *ctrA (T170A)* cells indeed proved the opposite phenotypes (Figure 3E, S5). For example, for the *divK* gene that negatively regulates CtrA activity and induces a G1 arrest when disrupted^11^, Tn insertions are 15-fold more frequent in *ctrA401* (T170I) cells versus *ctrA (T170A)* cells (Figure 3E, Supplementary Table S7).

### CtrA reprogramming in stationary phase is mediated by RpoB via (p)ppGpp

The fact that CtrA is implicated in the rickettsial growth phase (developmental) transition^18^ along with the role of reside T168 described above, prompted us to investigate the CtrA in stationary *C. crescentus* cells. We observed that the late S-phase promoter probe reporters P*_pleA_*-*lacZ* and P*_fliQ_*-*lacZ* are substantially upregulated in stationary phase compared to exponential phase *ctrA401* cells or compared to stationary *WT* cells (Figure 4A). The stationary phase promoter induction of these two promoters by CtrA401 and (WT) CtrA is enhanced by (p)ppGpp, the alarmone that is produced by SpoT in stationary phase^37^ as indicated by the fact the promoter activity is attenuated in Δ*spoT* cells. This finding prompted us to define the CtrA stationary phase target regulon by ChIP-Seq analysis in *WT* cells. We observed the ChIP-Seq profiles of CtrA to change substantially between exponential and stationary phase cells (Figure 4B, 4C). Moreover, robust CtrA occupancy on stationary phase chromatin shows that the presence of SpoT enhances chromatin occupancy of CtrA in stationary phase (Figure 4B) and that the activity of these CtrA-target promoters is indeed induced in stationary phase in a manner that depends on SpoT compared to exponential phase cells (Figure S6).

**Figure 4.**
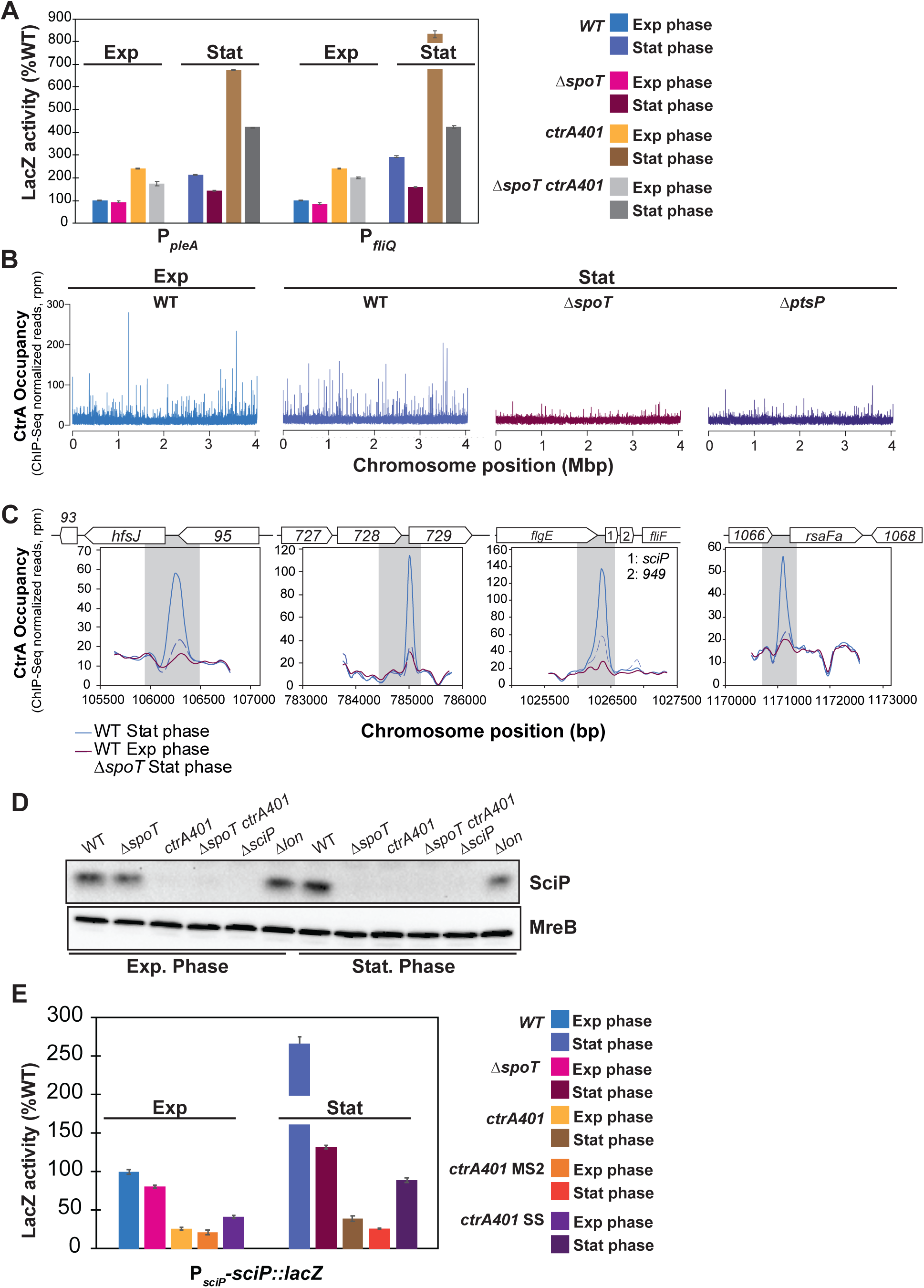
CtrA occupancy is substantially reprogrammed in stationary phase. A) Promoter-probe assays of transcriptional reporters carrying a *pleA* and *fliQ* promoters in *WT*, Δ*spoT*, *ctrA401* and derivatives. Transcription from P*pleA-lacZ* and P*fliQ-lacZ* in *WT* and *ctrA401* is strongly induced in stationary phase in a SpoT-dependent manner. Values are expressed as percentages (activity in *WT* in exponential phase is set at 100%). B) Genome wide occupancies of CtrA on the Caulobacter *WT* genome in exponential phase compared to CtrA occupancies in *WT*, Δ*spoT* and Δ*ptsP* genome in stationary phase as determined by ChIP-seq using antibodies to CtrA. The x-axis represents the nucleotide position on the genome (bp), whereas the y-axis shows the normalized ChIP profiles in read per million (rpm). CtrA occupancy is substantially reprogramed in stationary phase and is dependent on SpoT and PtsP. C) ChIP-seq traces of CtrA on different CtrA-binding promoter regions in *WT* cells in exponential phase, in *WT* and Δ*spoT* in stationary phase. Genes encoded are represented as boxes on the upper part of the graph, gene names and CCNA numbers gene annotation are indicated in the boxes or above. D) Immunoblot showing steady-state levels of SciP in *WT*, Δ*spoT*, *ctrA401*, Δ*spoT ctrA401*, Δ*sciP* and Δ*lon*, in exponential and stationary phase. MreB serves as a loading control. E) Promoter-probe assays of *sciP*-translational reporters in *WT*, Δ*spoT*, *ctrA401* and derivatives. Values are expressed as percentages (activity in *WT* set at 100%). Data from four independent experiments, error bars are standard deviation.

As further confirmation for the role of (p)ppGpp in enhancing CtrA promoter occupancy in stationary phase, ChIP-Seq also revealed that CtrA was also less abundant on chromatin isolated from stationary cells lacking PtsP, the E1 component of an alternate phosphoenolpyruvate transfer system that activates the SpoT pathway upon nitrogen starvation^38^ (Figure 4B). A prominent example of a stationary-phase induced CtrA-target promoter is the G1-phase promoter of the *hfsJ* gene (P*_hfsJ_*, see ChIP-Seq traces in Figure 4C). The strong SpoT-dependent increase in CtrA occupancy correlates well with the increase in HfsJ protein steady-state levels in stationary phase by immunoblotting (Figure 1B). CtrA steady-state levels in stationary phase cells are reduced in the absence of SpoT (Figure 5A, right), suggesting that occupancy of CtrA is enhanced by (p)ppGpp because it promotes an increase in CtrA abundance.

**Figure 5.**
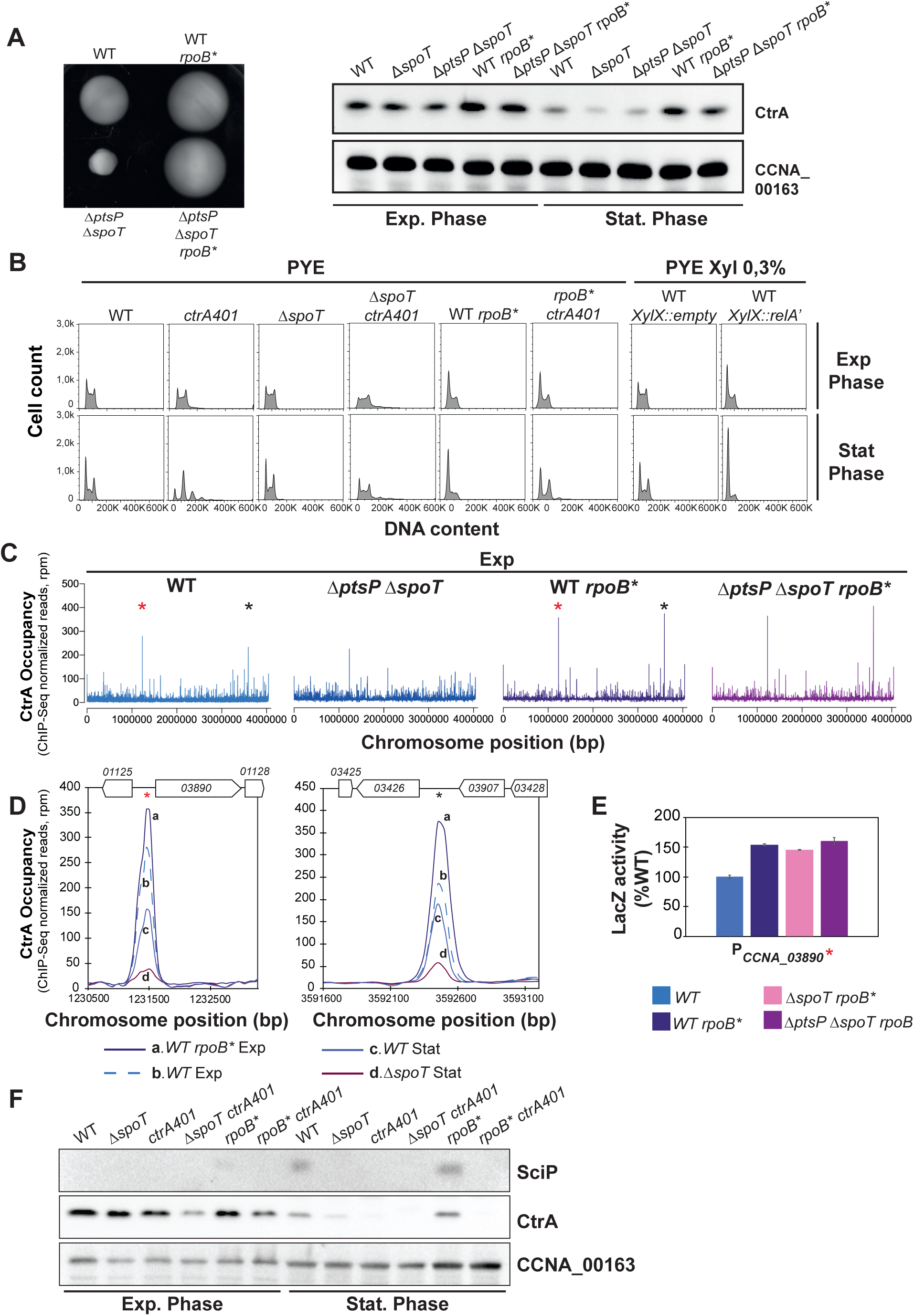
CtrA reprogramming in stationary phase is mediated by RpoB via (p)ppGpp. A) Identification of the *rpoB*^H559P^ mutation as motility suppressor of the Δ*ptsP* Δ*spoT* mutant that restores CtrA steady state levels in stationary phase. Swarm (0.3%) agar plates inoculated with *WT*, Δ*ptsP* Δ*spoT*, *WT rpoB\** and Δ*ptsP* Δ*spoT rpoB\**. *rpoB*^H559P^ mutation confers hypermotility to *WT* and Δ*ptsP* Δ*spoT* cells. Immunoblot analysis (right) of the steady-state levels of CtrA in exponential and stationary phase of *WT*, Δ*spoT*, Δ*ptsP* Δ*spoT*, *WT rpoB\** and Δ*ptsP* Δ*spoT rpoB\**. CCNA_00163 serves as a loading control. B) FACS analysis and DNA content (FL1-A) quantification in exponential and stationary phase of *WT*, Δ*spoT*, *ctrA401*, *rpoB\** and derivatives. Cells expressing the constitutive active form of *E. coli* RelA (RelA’) are also shown. *rpoB\** and ectopic induction of (p)ppGpp lead to G1 block in exponential and stationary phase. In stationary phase, *ctrA401* cells are elongated and do not replicate while disruption of *spoT* improves their phenotype. C) Genome wide occupancies of CtrA on the Caulobacter *WT*, Δ*ptsP* Δ*spoT*, *WT rpoB\** and Δ*ptsP* Δ*spoT rpoB\** genome in exponential phase. *rpoB\** mutation impacts the occupancy of CtrA on chromatin. Stars indicate 2 highly enriched peaks in *rpoB\** at the *CCNA_03890* and *CCNA_03426* loci. D) ChIP-seq traces of CtrA on the two CtrA-binding promoter regions highlighted with stars in C) in *WT* and *rpoB\** cells in exponential phase and in *WT* and Δ*spoT* cells in stationary phase. Genes encoded are represented as boxes on the upper part of the graph, gene names and CCNA numbers gene annotation are indicated in the boxes or above. E) Promoter-probe assays of transcriptional reporters carrying the *CCNA_03890* promoter in *WT*, *rpoB\**, Δ*spoT rpoB\** and Δ*ptsP* Δ*spoT rpoB\**. Values are expressed as percentages (activity in *WT* set at 100%). F) Immunoblots showing steady-state levels of SciP and CtrA in *WT*, Δ*spoT*, *ctrA401*, Δ*spoT ctrA401*, *rpoB\** and *rpoB* ctrA401* in exponential and stationary phase. CCNA_00163 serves as a loading control.

Interestingly, FACS analysis revealed a pronounced cell cycle defect of *ctrA401* cells in stationary phase compared to exponential phase or compared to stationary *WT* cells (Figure 5B). In fact, *ctrA401* cells accumulate extra chromosomes and there are few S-phase cells compared to *WT* populations. The accumulation of extra chromosomes in stationary *ctrA401* cells is mitigated by the Δ*spoT* mutation and the elevated (late S-phase) promoter activity in *ctrA401* cells is curbed when SpoT is inactivated (Table 2, Figure 4A). To test whether this contribution of SpoT/(p)ppGpp is mediated via allosteric control of RNA polymerase by (p)ppGpp, we sought suppressor mutations that render cells independent of (p)ppGpp for cell cycle control. To this end, we spotted Δ*spoT* Δ*ptsP* double mutant cells on swarm agar and isolated spontaneous suppressor mutants that overcome the swarming defect of the parent. Genome sequencing of one such suppressor mutant and backcrossing of the mutation revealed a change of histidine codon to a tyrosine at position 559 of the *rpoB* gene (*rpoB\**), encoding the beta-subunit of RNA polymerase, to confer the increase in swarming motility (Figure 5A left). The *rpoB\** mutation also augments swarming motility when introduced into *WT* cells (Figure 5A left) and increases the fraction of the G1 population as determined by FACS, akin to *WT* cells engineered to ectopically produce (p)ppGpp with an inducible heterologous (p)ppGpp synthetase gene from *E. coli*^39^ (*relA’*, Figure 5B). Consistent with these results, the steady-state levels of CtrA are also upregulated in cells with the *rpoB\** mutation (Figure 5A, right) and the ChIP-Seq profile of CtrA-bound promoters precipitated from chromatin of exponentially growing cells revealed that the *rpoB\** mutation impacts the distribution of CtrA on its targets *in vivo*. Several promoters that CtrA prominently binds to in *WT* exponential phase are further increased in *rpoB\** cells as confirmed by the ChIP-Seq traces presented in Figure 5D. The transcripts of *CCNA_03890* and *CCNA_03426*, the former encoding a hypothetical protein conserved in the *Caulobacterales* order and the latter a MarR-like transcription factor, are strongly cell-cycle regulated like other CtrA-regulated transcripts^40^, consistent with notion that their promoters are directly activated by CtrA and enhanced in the presence of (p)ppGpp. In exponential phase cells, the *rpoB\** mutation does not fully recapitulate the CtrA occupancy profile of stationary stationary phase *WT* cells (Figure 4B, 5C, 5D), indicating that unknown determinants other than (p)ppGpp play a role in CtrA reprograming at certain promoters upon the transition into stationary phase. Together the reprogramming of CtrA and strong increase in SciP expression in stationary phase in a manner that depends on (p)ppGpp (Figure 4D, 4E) provides further support that cell cycle transcriptional regulators also fulfill a role in growth phase transition signaled by (p)ppGpp.

## Discussion

### Coordination of G1-specific traits with CtrA

How CtrA can switch from activating late S-phase promoters to G1-phase promoters is fundamental to fully understand virulence gene expression underlying alpha-proteobacterial pathogenesis and symbiosis. In the human pathogen *Brucella abortus* G1-phase cells play an important role in driving the early stages of infection, while in the plant symbiont *Sinorhizobium meliloti* transcriptome studies have documented in synchronized populations the transcriptional switch from late S-phase to G1-phase^37, 41–43^. Upon studying the *C. crescentus ctrA401* mutant we discovered a determinant uncoupling the two promoter classes activation by CtrA. Importantly, we observed that binding of CtrA401 at late S-phase promoters is not compromised *in vivo*. Instead, it is elevated compared to *WT* with a commensurate increase in promoter activity, while G1-phase promoters show the inverse behavior.

In *C. crescentus*, the promoter switch drives cell surface structure remodeling, including pili and capsule that are known virulence determinants in bacterial pathogens^1^. The acquisition of pili in G1 cells is triggered by expression of the PilA pilin^44^, while capsule is lost from G1 cells upon expression of the negative regulator of capsulation, the HvyA transglutaminase homolog^5^. These traits collectively promote the susceptibility of G1 cells to the bacteriophages ϕCr30 and ϕCbK that bind the S-layer and pili, respectively^33, 34^. The *ctrA401* mutant is resistant to both phages because constitutive capsulation protects against infection by ϕCr30^5^, while loss of piliation is due to the defect in PilA expression in *ctrA401* cells and prevents infection by ϕCbK. Since cellular dispersal functions are activated by the CtrA dependent promoter switch, double stranded DNA phages such as ϕCr30 and ϕCbK may harness the opportunity of infecting G1 cells in order to introduce their genomes into the host before chromosome replication commences. Expression of HvyA not only reverses the constitutive capsulation phenotype of *ctrA401* cells, but also ameliorates the swarming motility defect, indicating that capsule interferes with flagellar motility, explaining why HvyA expression has been incorporated into the regulatory program of *Caulobacter* G1 cells and orthologs appear to underlie similar control. Thus, such coordination is likely critical for efficient dissemination and virulence properties of G1 cells in different alpha-proteobacteria.

### The pivotal role of residues T168 and T170 in CtrA

CtrA is essential and its predicted essentiality formed the basis for the isolation of a lethal “loss of function” (*ctrA401*) mutation that was isolated at the restrictive temperature of 37°C, while the second selection criterion was impaired negative regulation of class II flagellar (late S-phase) activated by CtrA^6^, despite the known role of CtrA being activating most target promoters^2^. How the negative and positive regulation by CtrA on the same class of promoters could be instated was an unresolved question. The results described here provide a rational explanation for this effect, as we showed that only activation of the G1-promoter class is curbed by the CtrA mutation. In principle, the T170I mutation in CtrA401 can also be viewed as a “gain of function” mutation endowing CtrA401 with an extended ChIP profile (Figure 1C), most likely reinforced by the reduction of *sciP* expression (Figure 1C, 1D; Figure S1C, S1D) at 30°C.

By contrast, activating function of CtrA (or stability) is lost at 37°C and thus cells suffer from insufficiency of essential cell division (and likely other) transcripts and/or de-regulation of DNA replication, ultimately causing cell death. Since the mutation in CtrA401 (T170I) lies in a highly conserved residue within the DNA-binding region, our findings unearth a critical residue directing promoter, gene expression and thus deterministic phenotypic switches. Molecular modelling suggests that the residues T170 and T168 do not engage in direct contacts with DNA, however they likely fulfill a critical role in proper positioning of the recognition helices or cooperative effects required for promoter binding and activation. It is possible that a small quantitative difference in affinity for the target box, especially in the context of architecture of G1 promoters, underlies or contributes to the qualitative block in the promoter switch of the CtrA401 mutant versus WT CtrA.

Our suppressors analyses revealed a compensatory mutation (T168I) in close proximity to the T170I mutation near the highly conserved DNA-binding region of CtrA (Figure 3B). This mutation not only partially reinstates the promoter switch to CtrA401 (Table 2), but also ameliorates the temperature sensitivity of *ctrA401* cells (Figure S4C). Homology-based structural predictions of CtrA401 do not suggest a modification of the DNA-binding domain structure compare to WT CtrA (Figure 3C). However, both mutations T168I and T170I lead to a reduction of at least two predicted β-sheets (β9- β10) that form a β-hairpin of the DNA-binding domain (DBD) and an increase in the length of the loop between them (Figure 3C) suggesting potential alteration of CtrA DNA-binding. Indeed, this C-terminal β-hairpin modified in the CtrA401-SS (T168I/T170I) double mutant likely forms hydrogen-bonds with the DNA-backbone as in the case of BasR/PmrA response regulator used to model CtrA structures^45^. Our modelling revealed several putative structure associated possibilities that could influence CtrA-dependent gene expression. While modelling of BasR/PmrA predicts a head-to-tail binding of the two DBDs within a dimer, T168 and T170 of CtrA are located close to important regions. In the predicted DBD structure number one (DBD-1, close to the receiver domain, Figure 3C), T168 is implicated in the interaction with the β7- β8 loop of the DBD-2, while T168 and T170 are very close to the β-hairpin in the DBD-2. Although P168 of PmrA has been assigned such a functional role from being within a hydrophobic cluster^45^, the residue corresponding to T170 in CtrA has not been implicated. Nevertheless, the T170 and T168 are very close and both may act on each other, for example to promote cooperativity on DNA-binding within certain promoter sequences or local DNA architectures imposed by other DNA-binding regulators. Interestingly, residue T168 of CtrA was previously identified as a gain-of-function mutation to improve activation of late S-phase promoters to Δ*mucR1/2* cells^12^ (Figure 1E). The difference between having a small structural change between an alanine and isoleucine instead of T168 is not impossible to reconcile with changes in promoter activation and cooperativity upon DNA-binding in these screens, but remains to be resolved at the atomic levels using structural studies. Interestingly, CtrA in the rickettsial lineage naturally has a valine in place of threonine at position 168 of *C. crescentus* CtrA. With our genetic implications of the isoleucine or alanine^12^ modification at this site in modulating the promoter switch of CtrA, depending on the context of a threonine or isoleucine at position 170, it will be revealing to investigate how rickettsial CtrA impact the suspect switch during the developmental cycle, especially since this lineage does not encode homologs of the accessory negative regulators, SciP and MucR, in their genomes^1^. Thus, we propose that functional Yin and Yang between T168 and T170 in *C. crescentus* CtrA fine-tunes the promoter switch by CtrA during the cell cycle and/or growth phase transitions.

### (p)ppGpp is required for promoter reprogramming by CtrA during growth transition

Another major change in CtrA promoter occupancy occurs upon entry in stationary phase and such reprogramming may reflect an ancestral role of CtrA that is still exploited by the obligate intracellular rickettsial pathogens to direct the growth phase transition of *Ehrlichia chaffeensis* from reticulate cells into dense-cored cells^18^. While *E. chaffeensis* does not offer a powerful genetic system to dissect how growth-phase dependent reprogramming of CtrA occurs, we show that in *C. crescentus* alarmone-based nutritional signals acting through RNAP play an important role. Having shown previously that P*_ctrA_* activity is increased in a (p)ppGpp dependent manner^46^, we confirmed that CtrA levels are decreased in stationary Δ*spoT* cells and that a mutation in RpoB (H559Y) bypasses the need for (p)ppGpp. This mutation also augments the steady-levels of CtrA, consistent with the previous evidence that (p)ppGpp regulates CtrA stability^46^. The role of (p)ppGpp in virulence and cell cycle control is well established *for B. abortus* and *S. meliloti*, respectively^37^. Moreover, suppressive mutations obtained in other alpha-proteobacteria that suppress (p)ppGpp null strain also mapped to *rpoB* and *rpoC*^47^. Interestingly, the conserved H559 residue in *C. crescentus* RpoB corresponds to the H551 of *Escherichia coli* that mimics (p)ppGpp binding^48^. Thus, the role of the alarmone in transcription regulation is conserved in bacteria, and its action on CtrA and its targets to implement promoter switching provides an ideal entry point to modulate cell cycle and growth phase transitions from a central regulatory node.

## Acknowledgements

Funding support is from Swiss National Science Foundatrion grant 31003A_182576 to PHV. We thank Mike Laub (Massachusetts Institute of Technology, USA), Lucy Shapiro (Stanford University, USA) and Justine Collier (University of Lausanne, CH) for materials, and Laurence Degeorges for excellent technical assistance.

## Author contributions

MD and PHV conceived and designed the experiments. MD, GP and PHV performed the experiments. All authors analysed the data. MD and PHV wrote the paper and all authors contributed to editing the manuscript.

