## Supplementary Figures and Material&Methods for "Bacterial cell cycle and growth phase switch by the essential transcriptional regulator CtrA"

### Figure Supplementary 1

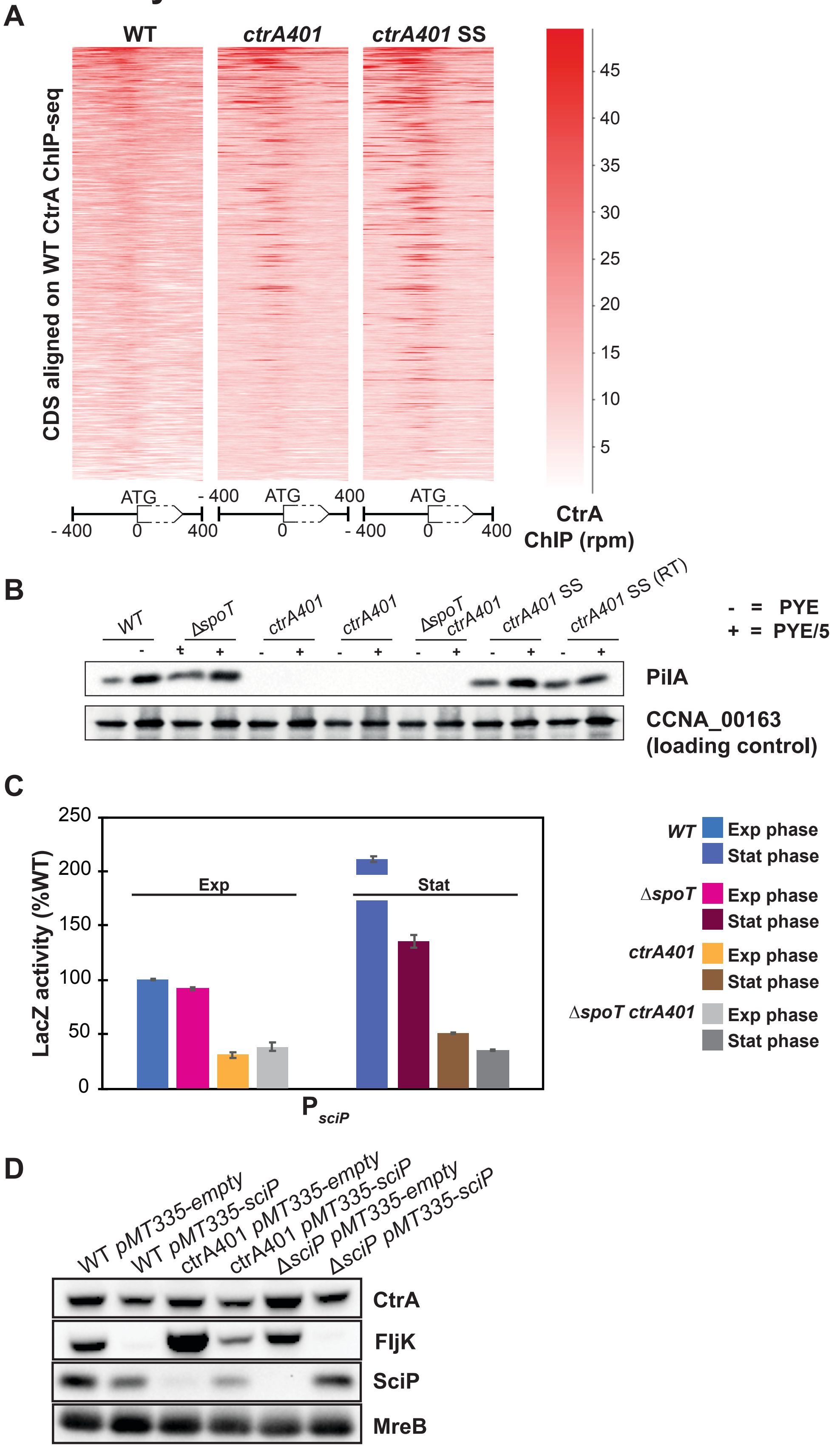


**Figure S1. Mutations in *ctrA* alter DNA-binding and G1-phase promoter expression.** A) Aligned probes plot of normalized CtrA, CtrA401 (T170I) and CtrA401-SS (T168I/T170I) chromosomal occupancy as determined by ChIP-seq using antibodies to CtrA. Caulobacter Coding DNA Sequence (CDS) are aligned over an 800 bp range around the translation initiation codon in *WT*, *ctrA401* and *ctrA401-SS* cells and sorted with reference to the relative abundance of reads compared to *WT*. CtrA401 (T170I) and CtrA401-SS (T168I/T170I) both show enrichment in occupancy on Caulobacter genome promoters. B) Immunoblots showing steady-state levels of PilA protein in *WT*, Δ*spoT*, *ctrA401* and *ctrA401-SS* in exponential phase cells grown in rich (PYE) or phosphate limited medium (PYE/5). *ctrA401* mutant cells do not accumulate PilA protein. CCNA_00163 serves as a loading control. C) Promoter-probe assays of transcriptional reporters carrying a *sciP* promoter in *WT*, Δ*spoT, ctrA401*, and Δ*spoT ctrA401* in exponential and stationary phase. Values are expressed as percentages (activity in *WT* set at 100%) D) Immunoblots showing steady-state levels of CtrA, FljK and SciP in *WT*, *ctrA401*, Δ*sciP* overexpressing SciP after 5 hours of growth in PYE supplemented with 50 µM Vanillate. The sciP overexpression plasmid *pMT335-sciP* leads to decrease in steady-state levels of SciP, CtrA and FljK while reduced levels of SciP in *ctrA401* or Δ*sciP* cells leads to an increase in steady-state levels of FljK. MreB serves as a loading control.

### Figure Supplementary 2

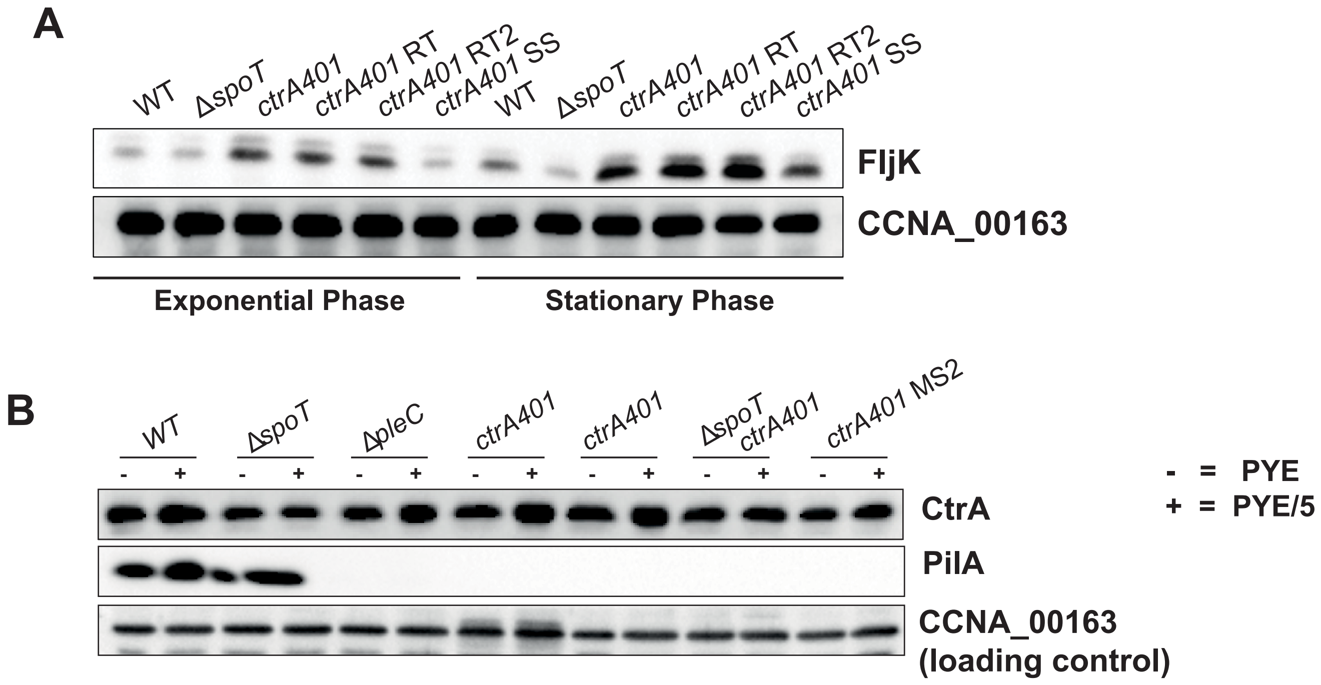


**Figure S2. Mutation in ctrA401 restores expression of CtrA targets**. A and B Immunoblotting experiments. A) Immunoblot showing steady state levels of FljK in *WT*, Δ*spoT*, *ctrA401* and *ctrA401-SS* in exponential and stationary phase growing cells. While *ctrA401* cells upregulate FljK steady-state levels, T168I mutation restores *WT* levels. B) Immunoblots showing steady-state levels of CtrA (and CtrA401) and PilA proteins in exponential *WT*, Δ*spoT*, Δ*pleC, ctrA401*, Δ*spoT ctrA401* and *ctrA401-MS2* cells grown in rich (PYE) or phosphate limited medium (PYE/5). CCNA_00163 serves as a loading control.

### Figure Supplementary 3

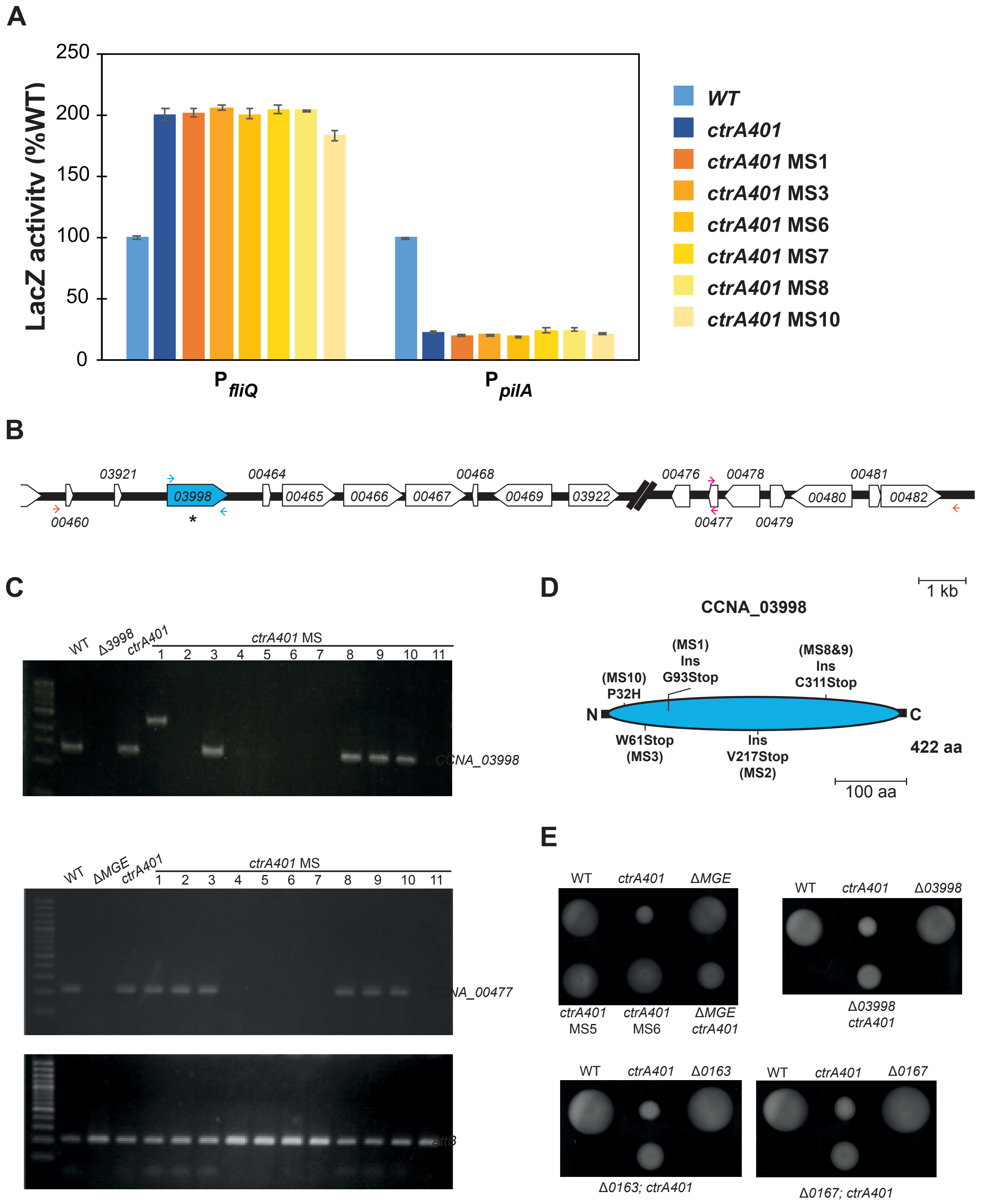


**Figure S3. Motility suppressors of *ctrA401* cells map to the *CCNA_03998* capsule biosynthesis gene.** A) Promoter-probe assays of transcriptional reporters carrying a *fliQ* and *pilA* promoters in *WT*, *ctrA401* and derivatives. Transcription from P*fliQ-lacZ* in *ctrA401* and *ctrA401-MS* is strongly compared to *WT* cells while transcription from P*pilA* is strongly reduced. Values are expressed as percentages (activity in *WT* is set at 100%). B) Schematic of the MGE locus identified through motility screen. Sequencing mapped SNP and IN/DEL mutations within the *CCNA_03998* for MS 1-3 and 8-10. B) Agarose (1%, upper) gel showing a PCR analysis of the *CCNA_03998* gene in the ten *ctrA401-MS* mutants. Primers forward and reverse were used to amplify the coding sequence. No amplification of *CCNA_03998* was obtained from *ctrA401-MS*4-7 cells similarly to the *ctrA401-MS*2. Agarose (2%) gels showing that no amplification of *CCNA_00477* (middle gel) was possible from *ctrA401-MS*4-7 cells. Primers forward and reverse were used to amplify the coding sequence of the *CCNA_00477* (upper gel). Primers forward and reverse were used to amplify the *attB* region (lower gel) showing the efficiency of excision of the MGE. D) Schematic of the CCNA_03998 protein. CCNA_03998 is a putative WbuB-like glycosyltransferase with a single domain (blue). Residues or IN/DEL identified in the motility screen are indicated. E) Swarm (0.3%) agar plates inoculated with *WT*, *ctrA401*, *ctrA401-MS*, ΔMGE, Δ *03998*. Δ*00163* and Δ*00167* and derivatives known to induce capsule-less phenotype are shown as control. Deletion of genes that confer the "heavy" (low buoyancy) phenotype by density gradient centrifugation improves the motility defect of *ctrA401* cells.

### Figure Supplementary 4

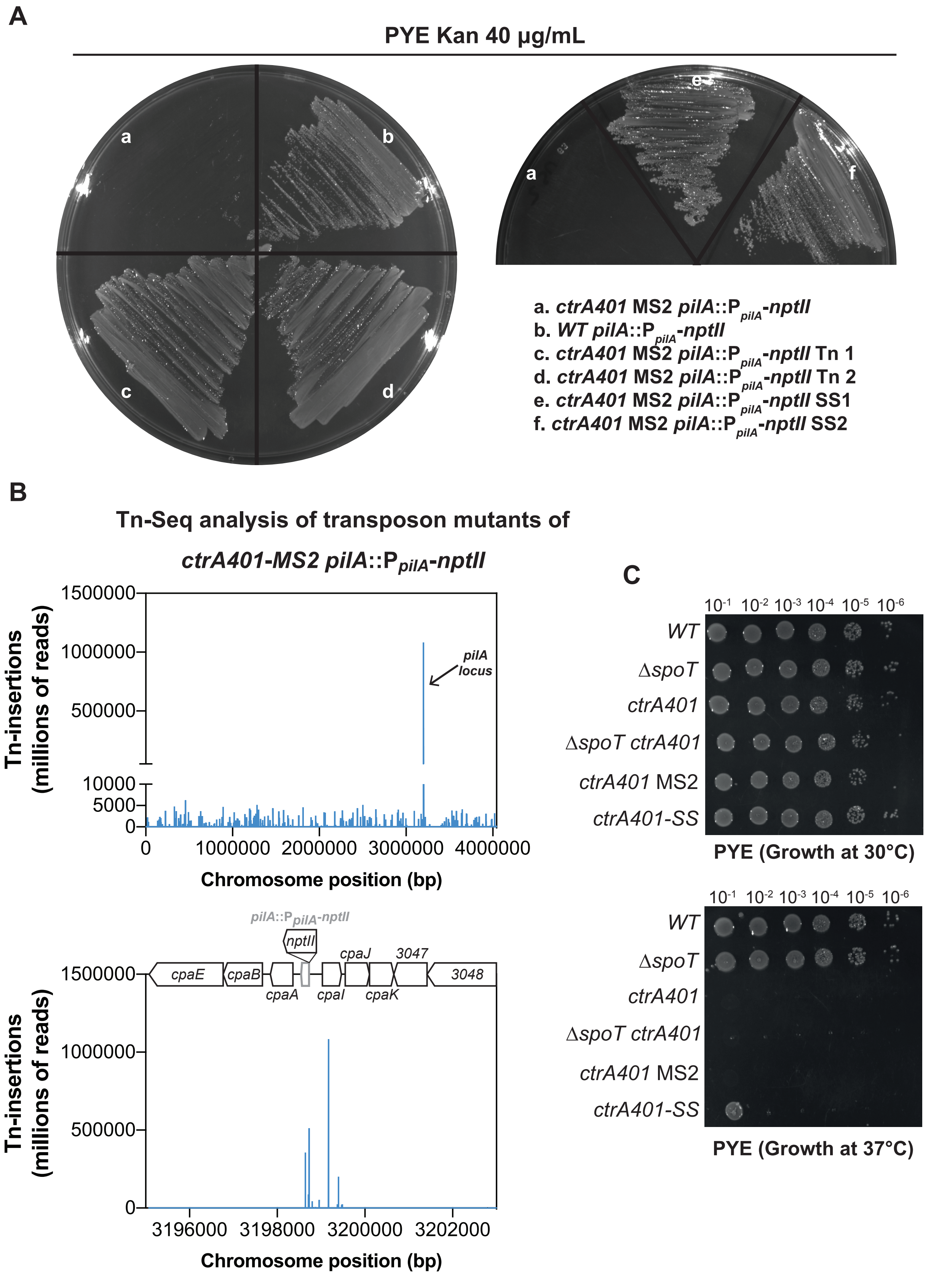


**Figure S4. Identification of the T168I mutation in *ctrA401* that restores activation of the P*pilA* promoter.** A) Growth of the *WT* (b), *ctrA401-MS*2 (a) and derivatives (c-f) carrying the *pilA::PpilA-nptII* transcriptional reporter on PYE plates containing 40 µg/mL of kanamycin. B) Graphs shown are genome-wide Tn-insertion profile of *ctrA401-MS2 pilA::PpilA-nptII* cells as determined by Tn-Seq (upper graph), as well as a zoom into the *pilA* locus where the Tn-insertions occurred (bottom graph). The abscissa shows the position as a function of the nucleotide coordinates, while the ordinate gives the corresponding abundance as Tn-insertion reads. C) Efficiency of plating assay of *WT*, *ctrA401* and *ctrA401-SS* cells on PYE agar plate at 30°C or 37°C. *ctrA401* and derivatives do not have loss of viability at 30°C. The T168I amino acid change slightly improves the thermo-sensitivity of the *ctrA401* mutant cells.

### Figure Supplementary 5

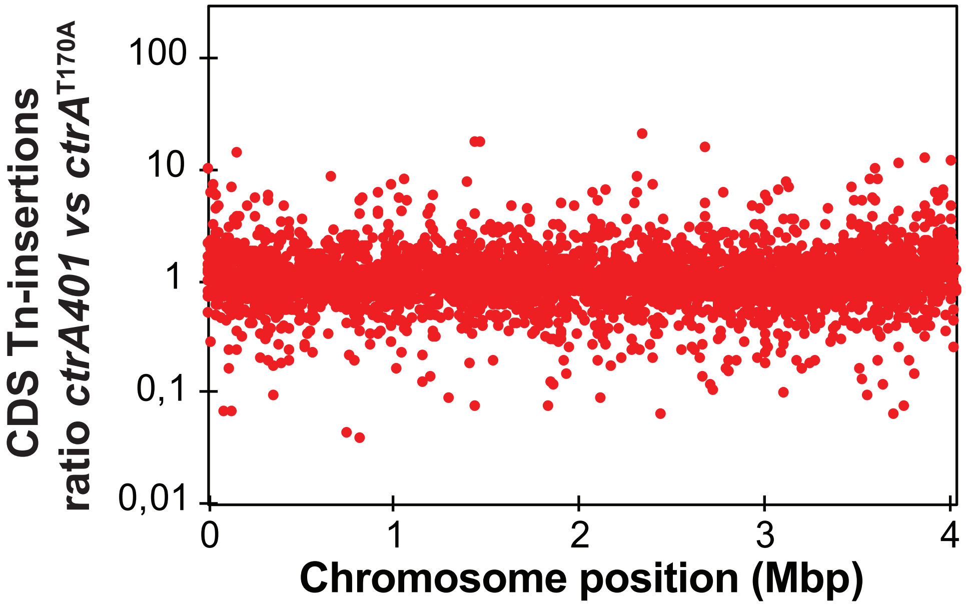


**Figure S6. Defining role of alanine or isoleucine at position 170 of CtrA.** Tn insertion bias in coding sequences (CDS) of *ctrA401* (T170I) cells relative to *ctrA*(T170A) cells as determined by Tn-Seq. The abscissa shows the position as function of nucleotide coordinates, and the ordinate provides the corresponding abundance as Tn-insertion reads.

### Figure Supplementary 6

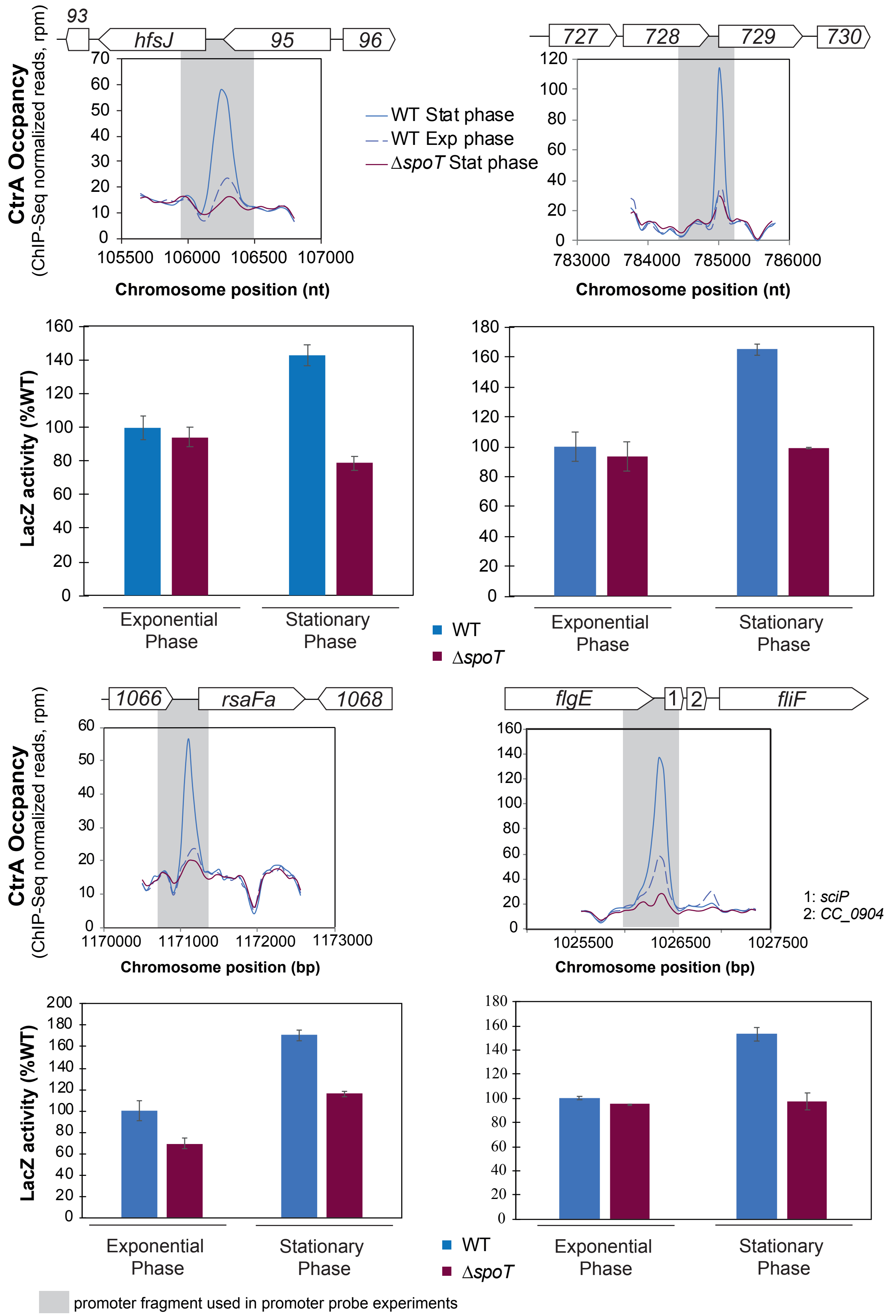


**Figure S5. Promoters occupancy of CtrA is reprogrammed in stationary phase in SpoT dependent manner.** ChIP-seq traces of CtrA in *WT* and Δ*spoT* mutant cells on the *hfsJ*, *CCNA_00729*, *rsaFa* and *sciP* promoters in exponential and stationary phase. Occupancy of CtrA is increased at the promoters tested in stationary phase and CtrA occupancy is reduced in Δ*spoT* mutant cells. Promoter-probe assays of transcriptional reporters in *WT* and Δ*spoT* cells in exponential and stationary phase in PYE confirmed the decrease in CtrA occupancy observed by ChIP-seq in the Δ*spoT* mutant cells in stationary phase. Values are expressed as percentages (activity in *WT* cells set at 100%).

**Table S1. DataStore summary report (excel file)**

**Table S2. ChIP-seq profiles of CtrA in *WT*, *ctrA401* (T170I) and *ctrA401-SS* (T168I/T170I) (excel file)**

**Table S3. ChIP-seq profiles of CtrA in *WT* cells in exponential and stationary phase, Δ*spoT* and Δ*ptsP* cells in stationary phase (excel file)**

**Table S4. ChIP-seq profiles of CtrA in *WT*, *rpoB**, Δ*ptsP* Δ*spoT* and Δ*ptsP* Δ*spoT rpoB** cells (excel file)**

**Table S5. Peak Calling (MACS2) analysis on CtrA ChIP-seq datasets (excel file)**

**Table S6. Tn-seq analysis of *ctrA401* MS2 *pilA::*$P_{pilA}$*-nptII* strain (excel file)**

**Table S7. Tn-Seq analysis of *WT*, *ctrA401* and *ctrA(T170I)* strains (excel file)**

### Supplementary Material and Methods

#### Bacterial Strains, Plasmids, and Oligonucleotides

#### Plasmid constructions

pNPTS138-*spoT*-us: PCR was used to amplify a DNA fragment upstream of the *CCNA_01622* or CC_1553 ORF. A 902 bp fragment (nt 1741409-1742310, NA1000 genome coordinates, flanked by an *Eco*RI site at the 5'end and a *Hind*III at the 3'end) was amplified using primers *spoT*-us-1-EcoRI and *spoT*-us-2-HindIII. The PCR fragment was digested with *Eco*RI/*Hind*III, then ligated into pNPTS138 restricted with *Eco*RI and *Hind*III.

pNPTS138-*rpoC-ds*: PCR was used to amplify a DNA fragment downstream of the *CCNA_00536* or CC_0502 ORF. A 873 bp fragment (nt 556213-557085, NA1000 genome coordinates, flanked by an *Eco*RI site at the 5'end and a *Hind*III at the 3'end) was amplified using primers *rpoC*-ds-1-EcoRI and *rpoC*-ds-2-HindIII. The PCR fragment was digested with *Eco*RI/*Hind*III, then ligated into pNPTS138 restricted with *Eco*RI and *Hind*III.

#### Strain constructions

Bi-parental intergeneric conjugations were used to transfer the resulting pNPTS138 derivatives into from *E. coli* to *C. crescentus* strains. The first plasmid integration event was selected for by resistance of the exconjugants to kanamycin and clones having undergone double recombination were obtained on PYE plates containing 3% sucrose. Mutants having undergone a double recombination that left the desired modification on the genome were identified by PCR using primers external to the DNA fragments used to make the pNPTS138 plasmids.

Construction of *WT* and Δ*ptsP* Δ*spoT* strains carrying *rpoB*$^{*H559P}$ mutation:

pNPTS138-*rpoC*-ds was first introduced into UG5591 (Δ*ptsP* Δ*spoT* strains carrying *rpoB*$^{*H559Y}$) by intergeneric conjugation and then plated on PYE harboring kanamycin (to select for recombinants) and nalidixic acid to counterselect *E. coli* donor cells. A single homologous recombination event close to the *CCNA_00536* locus of kanamycin-resistant colonies was verified by PCR. Second, bacteriophages ΦCr30 were prepared from UG5591 clones carrying the kanamycin resistance cassette and used to co-transduce the kanamycin-resistance cassette together with the *rpoB** $^{H559Y}$ point mutation into the wild type NA1000 and Δ*ptsP* Δ*spoT* the strain. The resulting strains were grown overnight in PYE medium lacking kanamycin. Cells were then plated on PYE supplemented with 3% sucrose and incubated at 30°C. Single colonies were picked and transferred in parallel onto plain PYE plates and PYE plates containing kanamycin. Kanamycin-sensitive cells, which had lost the integrated plasmid due to a second recombination event, leaving a point mutated version of *rpoB*, were then identified for mutation by PCR and Sanger sequencing.

The *ctrA401* derivatives were made from generalized transduction from the MD1343 (NA1000 *ctrA401* MS2 *ctrA::*pNPTS138-*ctrA-ds*) into *WT*, Δ*spoT*, ΔMGE, Δ*CCNA_00163*, Δ*CCNA_00167*, NA1000 *xylX*::pXTCYC4, NA1000 *xylX*::pXTCYC4-*relA'-M2*, NA1000 *rpoB**, Δ*spoT rpoB**.

Strains harbouring p*lacZ*-290 or derivatives were introduced into *C. crescentus* strains by electroporation and then plated on PYE containing tetracycline 1 µg/ml.

**Table S8. Plasmids used in this study**

| Plasmid | Relevant characteristics | Reference or source |
|---|---|---|
| pNPTS138 | Suicide vector used for gene replacement; Kana$^R$ | M.R.K. Alley unpublished |
| pNPTS138-*spoT*-us | pNPTS138 derivative carrying carrying a 902 bp fragment upstream of *spoT*; Kana$^R$ | This work |
| pNPTS138-*ctrA*-ds | pNPTS138 derivative carrying carrying a 920 bp fragment downstream of *ctrA*; Kana$^R$ | [1] |
| pNPTS138-*rpoC*-ds | pNPTS138 derivative carrying a 873 bp fragment downstream of *rpoC*; Kana$^R$ | This work |

| | | |
|---|---|---|
| pMT335 | Medium copy number plasmid for inducible expression; $P_{van}$, $Gm^R$ | 2 |
| pMT335-*TAP* | pMT335 derivative carrying the *TAP* fusion; $Gm^R$ | 3 |
| pMT335-*hvyA-TAP* | pMT335 derivative carrying the *hvyA-TAP* fusion; $Gm^R$ | 4 |
| pMT335-*sciP* | pMT335 derivative carrying the *sciP* ORF; $Gm^R$ | 1 |
| p*lacZ*290 | Low copy number vector containing *lacZ* gene (used to create *lacZ* transcriptional fusion); $Tet^R$ | 5 |
| p*lacZ*290-$P_{ctrA}$ | p*lacZ*290 containing *ctrA* promoter region; $Tet^R$ | 6 |
| p*lacZ*290-$P_{hvyA}$ (pSA205) | p*lacZ*290 containing *hvyA* promoter region; $Tet^R$ | 4 |
| p*lacZ*290-$P_{fliL}$ | p*lacZ*290 containing *fliL* promoter region; $Tet^R$ | 6 |
| p*lacZ*290-$P_{fljQ}$ | p*lacZ*290 containing *fljQ* promoter region; $Tet^R$ | 6 |
| p*lacZ*290-$P_{pilA}$ | p*lacZ*290 containing *pilA* promoter region; $Tet^R$ | 7 |
| P*lacZ*290-$P_{fliX}$ | p*lacZ*290 containing *fliX* promoter region; $Tet^R$ | 6 |
| p*lacZ*290-$P_{pleA}$ | p*lacZ*290 containing *pleA* promoter region; $Tet^R$ | 6 |
| p*lacZ*290-$P_{03790}$ | p*lacZ*290 containing *CC_3676* (*CCNA_03790*) promoter region; $Tet^R$ | 6 |
| p*lacZ*290-$P_{sciP}$ | p*lacZ*290 containing *sciP* promoter region; $Tet^R$ | 6 |

| | | |
|---|---|---|
| p*lacZ*290-$P_{ccrM}$ | p*lacZ*290 containing *ccrM* promoter region; $Tet^R$ | [8] |
| p*lacZ*290-$P_{spmX}$ | p*lacZ*290 containing *spmX* promoter region; $Tet^R$ | [9] |
| p*lacZ*290-$P_{hfsJ}$ | p*lacZ*290 containing *hfsJ* promoter region; $Tet^R$ | [6] |
| p*lacZ*290-$P_{2061}$ | p*lacZ*290 containing *CC_1982* (*CCNA_02061*) promoter region | [6] |
| p*lacZ*290-$P_{flaF}$ | p*lacZ*290 containing *flaF* promoter region; $Tet^R$ | [6] |
| P*lacZ*290-$P_{3890}$ | p*lacZ*290 containing *CCNA_03890* promoter region; $Tet^R$ | This work |
| p$P_{hvyA}$-*hvyA::lacZ* (pSA184) | pJC327 derivative carrying $P_{hvya}$-*hvyA::lacZ*; $Tet^R$; $Tet^R$ | [4] |
| pXTCYC4 | Non replicative vector in *C. crescentus* for expression of genes under the control of $P_{xyl}$ after the integration at chromosomal *xylX* locus; $Gm^R$ | [2] |
| pXTCYC4-*relA'*-FLAG | pXTCYC4 containing *relA'*-FLAG | [10] |

**Table S9. Strains used in this study**

| **Strains** | **Relevant characteristics** | **Reference or source** |
|---|---|---|
| ***Escherichia coli*** | | |
| S17 | RP4, Tc::Mu Km::Tn7 | [11] |

| | | |
|---|---|---|
| EC100D | *F- mcrA Δ(mrr-hsdRMS-mcrBC) Φ80dlacZΔM15 ΔlacX74 recA1 endA1 araD139 Δ(ara, leu)7697 galU galK λ- rpsL (StrR) nupG* | Epicentre |
| ***Caulobacter crescentus*** | | |
| NA1000 | Synchronizable variant of strain CB15 | [12] |
| LS3707 | NA1000 *ctrA401,* NA1000 derivative with the temperature-sensitive *ctrA401* allele | [13] |
| ML1749 | NA1000 Δ*sciP* ($tet^R$) | [14] |
| LS2382 | NA1000 Δ*lon* ($spc^R$) | [15] |
| FC769 | NA1000 Δ*spoT* | [16] |
| SS3 | Δ*ptsP* | [17] |
| LT1804 | Δ*ptsP* Δ*spoT* | This work |
| UG5591 | Δ*ptsP* Δ*spoT* motility suppressor (MS) ($rpoB^{H559Y}$) | This work |
| LT1805 | NA1000 *rpoB**$^{H559Y}$ | This work |
| LT1806 | Δ*ptsP* Δ*spoT rpoB**$^{H559Y}$ | This work |
| LT1807 | Δ*spoT rpoB**$^{H559Y}$ | This work |
| LT1808 | ΔMGE, NA1000 derivative with excision of the Mobile Genetic Element (MGE) | G. Panis (unpublished) |
| LT788 | Δ*CCNA_03998* | [4] |
| LT781 | Δ*CCNA_00163* | [4] |
| LT785 | Δ*CCNA_00167* | [4] |
| Δ*pleC* | NA1000 derivative with in-frame deletion of *pleC* | [18] |
| LT1809 | Δ*MGE ctrA401* | This work |
| LT1810 | Δ*CCNA_03998 ctrA401* | This work |
| LT1811 | Δ*CCNA_00163 ctrA401* | This work |
| LT1812 | Δ*CCNA_00167 ctrA401* | This work |
| LT1820 | NA1000 *ctrA401* MS1 | This work |
| LT1821 | NA1000 *ctrA401* MS2 | This work |
| LT1822 | NA1000 *ctrA401* MS3 | This work |
| LT1823 | NA1000 *ctrA401* MS4 | This work |
| LT1824 | NA1000 *ctrA401* MS5 | This work |
| LT1825 | NA1000 *ctrA401* MS6 | This work |
| LT1826 | NA1000 *ctrA401* MS7 | This work |

| | | |
|---|---|---|
| LT1827 | NA1000 *ctrA401* MS8 | This work |
| LT1828 | NA1000 *ctrA401* MS9 | This work |
| LT1829 | NA1000 *ctrA401* MS10 | This work |
| LT1831 | NA1000 *ctrA401* MS2 *pilA*::P$_{pilA}$-*nptII* | This work |
| LT1834 | NA1000 *ctrA401* MS2 *pilA*::P$_{pilA}$-*nptII* SS2 | This work |
| LT1845 | NA1000 *ctrA401* MS2 *ctrA::*pNPTS138-*ctrA-ds* | This work |
| LT1838 | NA1000 *ctrA*$^{T168I/T170I}$ (NA1000 *ctrA*-SS) | This work |
| LT1839 | Δ*spoT ctrA401* | This work |
| LT1840 | Δ*spoT ctrA*$^{T168I/T170I}$ | This work |
| JC835 | NA1000 *xylX*::pXTCYC4 | [10] |
| JC820 | NA1000 *xylX*::pXTCYC4-*relA'-M2* | [10] |
| LT1841 | NA1000 *xylX*::pXTCYC4 *ctrA401* | This work |
| LT1842 | NA1000 *xylX*::pXTCYC4-*relA'-M2 ctrA401* | This work |
| LT1843 | NA1000 *rpoB**$^{H559Y}$ *ctrA401* | This work |
| LT1844 | Δ*spoT rpoB**$^{H559Y}$ *ctrA401* | This work |

**Table S10. Oligonucleotides used in this study**

| Primers | Sequence 5'-3' |
|---|---|
| *spoT*-us-1-EcoRI | AAAAAAGAATTCCGCGCCCACCAGCGGCTGTTTGT |
| *spoT*-us-2-HindIII | AAAAAAAAGCTTAGCCGTGCATGCGCATCGCATAGACA |
| *rpoC*-ds-1-EcoRI | AAAAAAGAATTCAAACAGTCGAAGACCCGGACGCCAAAA |
| *rpoC*-ds-2-hindIII | AAAAAAAAGCTTCCCATATAGCCGGTCATGCCGAGCA |
| *attB*-FW | TCGGTCAGCACGACACAGATT |
| *attB*-RV | TCGACCTCGGGCGGCAATT |
| *477*-NdeI | AAAAAACATATGACCGAAGATCCATCCGACCT |
| *477*-EcoRI | AAAAAAGAATTCGATTGATGGCGTTAC |
| *3998*_NdeI | AAAAAACATATGCCAGACAAACGTATAAAGCTT |
| *3998*_EcoRI | AAAAAAGAATTCTAGGCGCACACGCGCTTCA |
| ctrA_NdeI | AAAAAAAAACATATGCGCGTACTGTTGATCGA |
| ctrA_XbaI | AAAAAATCTAGATGGGTGGGTTCTACAGATACGCT |
| rpoB-seq1 | AAAAAAATGCCGCACGACCTGATCAA |

| | |
|---|---|
| rpoB-seq2 | AAAAAATTTTGCAGGAGGGTCGGTT |
| P3890-EcoRI | AAAAAAGAATTCCTGAGCGCTTTCTAGCCTGCTT |
| P3890-XbaI | AAAAAATCTAGAACCTCGCGCCAAGCCCCCTTGAA |
